# An Information-Theoretic Approach to Reward Rate Optimization in the Tradeoff Between Controlled and Automatic Processing in Neural Network Architectures

**DOI:** 10.1101/2023.09.18.558214

**Authors:** Giovanni Petri, Sebastian Musslick, Jonathan D. Cohen

## Abstract

This article introduces a quantitative approach to modeling the cost of control in a neural network architecture when it is required to execute one or more simultaneous tasks, and its relationship to automaticity. We begin by formalizing two forms of cost associated with a given level of performance: an *intensity cost* that quantifies how much information must be added to the input to achieve the desired response for a given task, that we treat as the contribution of *control* ; and an *interaction cost* that quantifies the degree to which performance is degraded as a result of interference between processes responsible for performing two or more tasks, that we treat as inversely related to *automaticity*. We develop a formal expression of the relationship between these two costs, and use this to derive the optimal control policy for a desired level of performance. We use that, in turn, to quantify the tradeoff between control and automaticity, and suggest how this can be used as a normative framework for understanding how people adjudicate between the benefits of control and automaticity.

## I. INTRODUCTION

### Control-dependent processing versus automaticity

One of the most striking features of human cognition is the ability to adapt behavior over both short and long terms. The former confers remarkable flexibility in responding to novel environments, while the latter can lead to considerable improvements in the efficacy of processing in stable environments. Rapid adjustments of behavior to novel circumstances rely on the capacity for cognitive control, and are a fundamental component of characteristically human capabilities, such as problem-solving, planning, and language [1]. While this ability allows remarkable flexibility, it is generally associated with limitations in processing. The most salient of these are: a strict constraint on the number of control-dependent tasks that can be performed simultaneously (e.g., the ability to talk with a passenger when first learning to drive a car, or to sing while learning to play a musical instrument [2– 4]); and processing costs associated with switching from one task to another (e.g., when switching from parsing an equation to talking to a friend [5, 6]). In many cases these limitations can be overcome with sufficient learning, and tasks can come to be performed more effectively and, in some cases, even simultaneously with others [7–9]. However, this requires an investment of practice, and the resulting improvements in processing efficacy are largely tailored to that particular task (e.g., learning to play one instrument does not transfer fully to others [10]). This long-term adaptation is often referred to as the acquisition of automaticity for a task [7, 11].

The distinction between control-dependent and automatic processing is one of the cornerstones of cognitive psychology, and ample progress has been made in understanding the mechanisms that underlie this distinction [1, 12]. Considerable effort has been made to characterize computational mechanisms that can explain behavioral observations, and in some cases how these may be implemented in the brain. Recently, formal analyses have been proposed of some of the factors relevant to the allocation of control [13–19], and to the acquisition of automaticity [20]. Most recently, these analyses have begun to address the tradeoff between control and automaticity in neural network architectures—a tradeoff that is fundamental to understanding the adaptive nature of human behavior, and may be instructive in designing artificial systems with capabilities similar to those of humans [21]. Here, we build on this work, to develop a formally rigorous, normative framework for quantifying the costs associated with control-dependent processing versus automaticity, and the tradeoff between these.

### An information-theoretic approach

Following previous efforts [13, 14, 18, 19], we cast automaticity and control in information-theoretic terms: as the probability, given one or more stimuli (information), of encoding these and generating a task-appropriate response to each (i.e., executing the relevant actions, whether internal, such as encoding in longer term memory, or external, such as overtly motor responses). We use this approach, in relation to neural network processing architectures, to evaluate the processing efficacy of a network in executing a set of tasks, by encoding a set of stimuli as a pattern of activity over sets of input units corresponding to the stimulus dimensions relevant to those tasks, and propagating those inputs over one or more layers of intermediate (“hidden”) processing units, to generate a pattern of activity over a set of output units that determine the probability of response for each task. Implemented in neural networks, this approach has been used widely to account for human performance and underlying neural mechanisms in a wide range of cognitive paradigms [21–24] and, in machine learning, to build systems that approximate or exceed human performance in many of those tasks and others [25, 26].

### A fundamental tradeoff between control-dependent and automatic processing

Neural network formalisms have also been used within cognitive psychology and neuroscience to address the mechanisms responsible for control-dependent processing and automaticity, and the fundamental computational tradeoff between these [12]. However, these efforts have largely focused on relatively simple networks that are tractable to exhaustive numerical analysis (an example of which we provide in the following sections). In this article, we build on these efforts, and bring them into contact with information-theoretic approaches, to address the tradeoffs between control and automaticity both from a normative perspective and in more complex, multitask networks. In this section, we outline the key assumptions of the approach, drawn from the neural network literature.

First, we assume that the allocation of control is implemented as a set of biases that are used to regulate the activity of units required for executing a given task, both to insure that there is sufficient activity to overcome a response threshold for that task (increasing its probability of execution), and to compete effectively with other tasks that share processing units with the task by preventing them from activating conflicting representations in those units (thereby increasing the probability of *accurate* execution [12, 21, 27, 28]). Adjustments of these biases in the service of control are assumed to operate over short-time scales, thus enabling the flexible switching from one task to another [12, 29–3 Furthermore, to the extent that control confers the ability to use shared, general-purpose representations for different tasks, it affords the flexibility to perform novel tasks—a hallmark of the capacity for cognitive control [21]. However, this comes at a cost: the need to serialize processing in order to avoid the potential for conflict posed by the use of shared representations for different purposes at the *same time* [28]. This seriality constraint, that comes with the flexibility of processing, is also a hallmark of cognitive control. Together, they reflect a fundamental tradeoff in network architectures posed by the use of shared representations and the attendant requirement for control: they facilitate learning and generalization, but compromise the efficiency of processing afforded parallel execution [12]. Recent work has begun to formalize this tradeoff [17, 19, 21, 32], the broader implications of which we will consider in the Discussion. Here, we build on this idea, by formalizing the costs associated with the use of control, and addressing the implications it has for the choice between reliance on control-dependent processing versus the development of automaticity.

In contrast to control-dependent processing, we assume that automaticity is associated with the adjustment of connection weights that occurs over a longer time scale through training [27, 33]. This can strengthen an existing processing pathway responsible for executing a task, allowing it to more readily overcome a response threshold, compete more effectively with other tasks with which it shares representations, or induce a new pathway using separate processing units that are dedicated to that task and that isolate it from others, thus diminishing the likelihood of conflict with or interference from them and the attendant dependence on control [20, 21]. This can also be exploited to augment processing efficiency, through parallel task execution. However, the development of automaticity requires training, involving gradual, incremental adjustment of weights to insure a stable statistical encoding of the relevant associations [34, 35]. Thus, in addition to conferring the potential for greater efficiency of processing, automaticity also confers a form of flexibility, but over a longer time frame and with a different set of attendant costs than does control.

### Benefits and costs

As outlined above, both control-dependent processing and automaticity have their benefits and costs. In previous work, this tradeoff has been explored in the context of specific tasks, with respect to both the speed and accuracy of processing [27, 28, 36, 37]. However, while these efforts have been useful in specifying computationally explicit mechanisms underlying automaticity and control, they have focused on networks configured to perform a small number of tasks (usually just 2 or 3), and for the most part, have not addressed the question of whether or how the system optimizes the balance between automaticity and dependence on control— that is, they have not provided a normative account of this tradeoff. In this article, we develop an information theoretic approach to addressing these challenges. We do so by casting benefits in terms of the amount of reward accrued—proportional to the accuracy of processing— per unit time; that is, *expected reward rate*. Accordingly, costs can arise in either of two ways: degradation in processing accuracy, or prolongation of execution. Theoretical work in cognitive science [38] and neural network modeling [21, 27, 37, 39, 40] suggests that these may be different expressions of a single underlying factor—strength of processing—interacting with strategic choices regarding control (e.g., by regulating speed-accuracy tradeoffs and/or the allocation of attention). These align with the formulations of control and automaticity described above: with strength of processing determined by the connection weights in the processing pathway required to execute a specified task, and control determined by the biases allocated to the processing units responsible for task execution. Previous work within this framework, both theoretical and empirical, has shown how control parameters can be optimized to maximize reward rate. For example, for a given set of connection weights in a simple one-layered network, the speed-accuracy tradeoff is determined by the threshold of activity on the output layer (that is used to determine an overt response); there is a unique such threshold that optimizes reward rate; and, in many instances, this has been shown to accurately describe human performance in correspondingly simple decision making tasks [37, 41–4 Here, we assume that control can be used to achieve such threshold optimization by applying the appropriate bias to the units responsible for implementing the decision process (e.g., [45]), thus ensuring the optimal balance of speed and accuracy for a given strength of processing. Similarly, in settings requiring switching between tasks, the amount of control allocated to each can be optimized to balance the tradeoff between performance of each and the costs associated with switching between them, in order to maximize overall reward rate [36, 46, 47]. Thus, the benefits and costs of processing can be optimized with respect to reward rate.

The optimization of reward rate can be considered with respect to benefits and costs that accrue over a range of timescales of task performance, from the level of individual instances of performance (e.g., a “trial” of an experimental task) and/or the transition between these, to extended sequences of trials, and even longer timescales associated with training and practice. Neural network (and closely related dynamical systems) analyses of the optimization of the speed-accuracy tradeoff and/or task switching at the single trial (and trial-to-trial) timescale have provided valuable insights into the basic mechanisms underlying cognitive control, and how it may be implemented in neural architectures [29, 30, 46, 48]. While the timescale of the effects for an individual trial (and transitions between them) is generally small (on the order of hundreds of milliseconds), optimization can have a substantial impact on reward rate; for example, when the stakes for performance are very high, or over repeated performance of the same task(s). Nevertheless, the timescale of these effects contrasts with two other forms of temporal costs that can be an order of magnitude larger, and that are the focus of this article. These concern the serialization of the performance of multiple tasks forced by the reliance on shared representations (and enforced by control), and the training time required to acquire separated, task-dedicated representations that support parallel processing (i.e., to acquire automaticity). We briefly consider each of these below.

### Serialization costs

The use of shared representations confers a benefit on performance by allowing new tasks to be acquired quickly. However, as suggested above, if more than one of the tasks that share those representations is to be performed, this carries a temporal cost imposed by the additional time required to serialize execution (through the allocation of control), as compared to their simultaneous parallel execution when they rely on separate, task-dedicated representations. We refer to this as the *serialization cost*, and assume this is proportional to the total number of tasks to be executed [49, 50]. We should note that the serialization cost of control-dependent processing is closely associated with a corollary temporal cost of switching from one task to another, commonly referred to as the *switch cost* [5, 6, 21, 51], that is, in addition to the time spent on performing each task individually. However, as noted above, switch costs are generally an order of magnitude smaller than the serialization cost itself, and thus, for simplicity, they are not included in the analyses presented in this article. In the Discussion (Section V), we consider how the framework might be extended to include these in the future.

### Learning costs

As opposed to control-dependent processing, the benefits of automaticity are not only more rapid performance of each individual task but, critically, the potential for parallel execution of multiple tasks [7]. The efficiency that comes from multitasking obviates the serialization cost attached to control-dependent processing. This stems from reliance on more specialized, or fully task-dedicated representations, which makes them less likely to conflict with others, and therefore more amenable to execution in parallel [8, 21, 52–54]. However, these benefits also carry a temporal cost, in this case, the (often considerable) amount of time required to achieve automaticity through practice [7, 9, 55]. We refer to this as the *learning cost*. Thus, while control-dependent processing is associated with seriality (and switch costs), automaticity is associated with a learning cost.

### An intertemporal choice

The different types of cost associated with control-dependent processing and automaticity pose a fundamental intertemporal choice–and corresponding optimization problem–with regard to reward rate maximization: Under what circumstances should the more immediate benefits of control-dependent processing (as a consequence of rapid task acquisition) be favored, at the expense of less efficient serial processing, as compared to the benefits of automaticity (more efficient, parallel processing) that come at the expense of greater time required for acquisition [49, 50]. This tradeoff is familiar to anyone who has considered the relative advantages of hunt-and-peck typing versus learning to touch type, and similarly for using the keyboard of a musical instrument. It is also likely to be relevant to the ability of autonomous artificial agents to adapt and optimize their behavior in changing environments (to which we return in the Discussion).

To formalize and analyze this problem, in this article we quantify the costs of control-dependent versus automatic processing at two levels of description, the *neural network* level and the *task graph* level, that are closely related to one another. We include the former, as neural networks have been useful in providing well-specified process models of control and automaticity [21, 27, 29]. Such models implement tasks at the scale of processing units used to represent individual stimuli, responses, and often internal (“hidden”) units used to associate stimuli to responses.[122] Stimulus and response units are often divided into subsets of units (e.g., corresponding to different stimulus and response dimensions, such as colors versus shapes and verbal versus manual responses), with tasks implemented as pathways that map a set of stimulus representations to a set of response representations, possibly by way of one or more layers of hidden representations (Figure 1, left panel). Such models have been used to explain detailed patterns of performance in a wide range of tasks that address both the effects of different stimulus conditions and the dynamics of processing. However, such models are often intractable to closed-form analysis, and become even more of a challenge when used to address how interactions among multiple tasks affect performance. Accordingly, we address interactions among tasks at the level of a task graph. This builds on a simplification of the neural network implementation of an individual task, which summarizes its relevant properties (under assumptions we describe below), and allows us to formally analyze the interaction among multiple tasks. In a task graph, each node corresponds to one of the sets of processing units in a neural network model that represent a particular type of information (e.g., a stimulus dimension such as colors or shapes, or a particular type of response such as verbal or manual (Figure 1, right panel), and each edge corresponds to the weights of the connections from the set of processing units represented by one node to the processing units represented by another. Thus, the edge connecting an input node (representing a particular stimulus dimension) to an output node (representing a type of response) corresponds to the pathway in a neural network model that maps the set of stimulus units to the set of response units used to perform a given task. Following prior information-theoretic formalisms [13, 18], we use the task graph to define the costs of automaticity and control with respect to the formation and use of such mappings.

**Figure 1.**
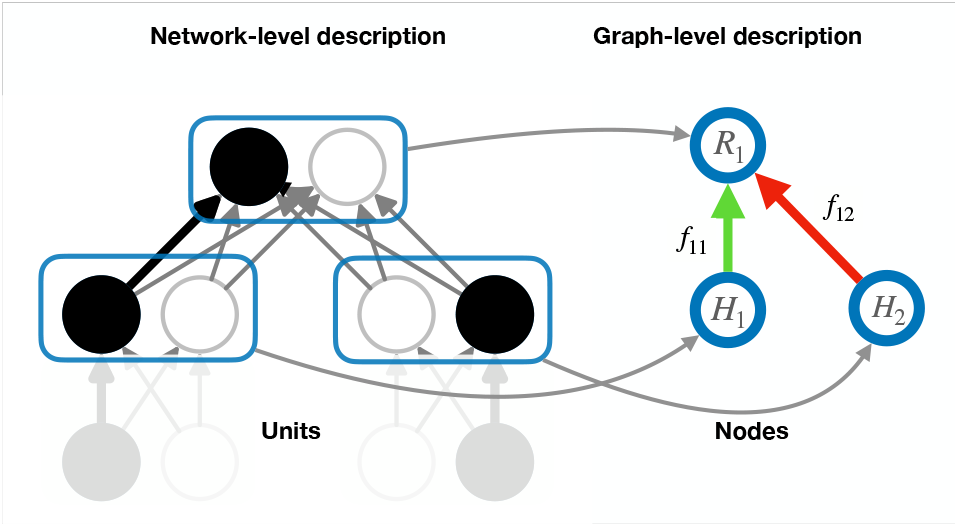
Correspondence of neural network and task graph representations of two tasks. *Neural network level:* shows the processing units in a neural network that implement two tasks, each of which maps the two stimulus units in each of two separate stimulus sets to the two response units of a shared response set. *Graph:* shows the corresponding task graph, in which each node corresponds to the processing units in a given set (outlined in blue) in the neural network, and each edge (red and green arrows) represents the set of connections (shown in gray) between the sets of processing units represented by the corresponding nodes. Note that in this task graph, we illustrate only the mapping from internal (“hidden”) representations corresponding to two sources of stimulus information to the set of response representations that are shared by them. For a more complete treatment that includes the relationship of external stimuli to their internal representations, see [57].

Beginning with single tasks (along the lines of [13]), we extend the formalism to address an arbitrary number of tasks and consider how performance can be optimized when confronted with the requirement to perform two or more of them. We investigate whether the benefits are greatest from the use of shared representations and control (at the expense of serialization) versus separated representations and automatic parallel processing (at the expense of additional training). We do so by defining a metric that specifies the expected level of interference between processes for a given network configuration, and use that to derive a corresponding metric for the amount of control (in terms of biasing parameters) required to mitigate this interference for a desired level of expected processing efficacy. We then discuss how this metric can be integrated with a corresponding metric for the learning cost of automaticity, that specifies the extent to which a network architecture must be adjusted through weight modifications (i.e., the formation of separated, task-dedicated representations) to mitigate multitasking interference and support parallelization of performance. In each case, we quantify the benefits and costs in terms of the common currency of time, thus permitting a normative analysis of the tradeoff between reliance on control-dependent processing versus the acquisition of automaticity.

## II. AN INFORMATION-THEORETIC FORMULATION OF AUTOMATICITY AND CONTROL

### A. Single Task Processing: Reflexes, Automaticity, and Control

To consider control-dependent processing within an information theoretic framework, Koechlin and Summer-field [13] proposed an expression for probability of performing a single task given a relevant stimulus. More precisely, to select a given action *a* with relative frequency *p*(*a*), they used standard Shannon entropy as a measure of the information required to select the action *a* irrespective of the stimulus, i.e.

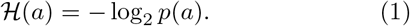

We refer to this as the *information cost* of action a. However, this formulation does not explicitly take into account the distribution of stimuli in an environment (i.e. the relative frequency of encountering a given stimulus or set of stimuli), which can be done by considering the mutual information ℐ (*s, a*) between a stimulus s and the associated action a, written as:

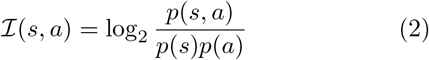

This, in turn, leads to the natural expression for the amount of information needed to select action *a* given the stimulus *s*:

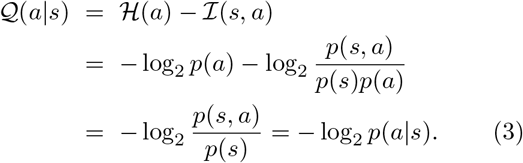

Here 𝒬 measures the extra information contained in observing the action a given that stimulus s was present (ℐ; (*s, a*)), compared with the case in which no information about the stimulus is available (ℋ (*a*)). Note that the functional forms of 𝒬 (*a* |*s*) and ℋ (*a*) are the same, but they are given different probability distributions (*p*(*a* | *s*) and *p*(*a*), respectively; see [18] for a related formulation).

#### Reflexes

Given the formulation above, 𝒬 (*a* | *s*) effectively captures the degree to which an action is selected irrespective of current task goal (i.e., context): if action a is *always* executed when a stimulus s is present (i.e. *p*(a | *s*) = 1), then 𝒬 (*a* | *s*) = 0. We define such actions as *reflexes*, insofar as they occur independently of task context or the allocation of control, and treat them as an extreme form of automaticity. It is important to note, here, that whereas the term “actions” connotes overt motor responses, the formulation applies equally well to covert, internal actions, such as the encoding of a stimulus into memory or the elicitation of emotional response, that do not have immediate observable consequences.

#### Automaticity

While reflexes that elicit overt actions can be thought of as a limiting case of automaticity, the latter more commonly refers to processes that, although they do not elicit an overt action, and have not been allocated control, nevertheless, elicit internal responses that are strong enough to interfere with competing processes to which control *has* been allocated. Word reading is widely regarded as a canonical example of this [11, 58]. This is evidenced by the classic Stroop effect [27, 59, 60], in which the response for naming the color in which a word is printed is slowed for incongruent stimuli (e.g., the stimulus GREEN)— that is, when the word itself is different than the color, and thus conveys competing information. This is taken as evidence of the automaticity of word reading, insofar as the effect occurs despite the instructions and intent to ignore the orthographic features of the stimulus. That is, processing of the word appears to occur (at least to some extent) independently of control, and thereby interfere with the processes required for color naming, to which control *has* been allocated, even when no overt response is made to the word. According to the formulation above, the internal processes of encoding the orthographic features of the stimulus, and associating these with phonological representations used to generate verbal responses, might be thought of as reflexive. Nevertheless, these processes are not *so strong* as to command an obligate overt response to the word or fully corrupt a response to the color: When confronted with a written word, people do not reflexively blurt these out loud without instructions (or intent) to do so; and, in the Stroop paradigm, people are able to name the color of incongruent stimuli with near perfect accuracy (i.e., degradation in performance manifests primarily in slower response time, not erroneous responses). Furthermore, there is considerable empirical evidence that word reading is sensitive in some degree to attentional demands (e.g., [61, 62]). Thus, word reading is an example of a task that is considered to be automatic, insofar as it engages internal processes that are strong enough to interfere with others that rely on similar representations (e.g., the phonological codes required for naming colors), but not so strong as to obligatorily elicit an overt action or to be fully independent of the effects of control. That is, overt responding *always* requires some additional support from control.

#### Control

While the foregoing suggests that tasks that are not overtly reflexive require *some* degree of support from control (e.g., word reading), that support must be augmented if the overt response to a stimulus must compete with a different one favored by a stronger, competing task (e.g., color naming). This suggests that a task’s dependence on control (or, conversely, its degree of automaticity) should be viewed both as a *continuous* attribute that is determined by the strength of the processing pathway required to execute the task, and as a *relative* attribute that is determined by any other tasks with which it may be in competition (see [27] for a more elaborate treatment of this point). These two attributes align with the two biasing effects of control introduced above—allowing a process to exceed a response threshold and/or diminish interference from competing processes— that are central to the formal treatment of its relationship to automaticity we present below. Critically, they suggest that, other than for reflexes, performance relies on the *context* in which a task is executed, which includes both the stimuli present in the environment (that may elicit other competing tasks) as well as internal signals (e.g., task goals) used for control.

For cases in which a response to a particular stimulus depends on the context in which it occurs, Koechling and Summerfield proposed to add terms to the expression for 𝒬 (*a* | *s*) that are sensitive to the context, *c*, in which a stimulus appears [13]. In the same way as Eq. 3, this splits the expression as follows:

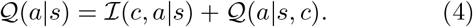

Here the first term refers to the additional information encoded by the context in presence of the stimulus *s*, and the second to past experiences. Importantly, context can be thought of as determining the task that should be performed (i.e., the stimulus-response mapping that is appropriate for that setting). To emphasize this point, and to align with the formulation of control as a signal that provides an internal representation of context used to specify which *task* should be performed, we replace the term c with t. Accordingly, the same pair (a, s) can have different probabilities, and hence informational demands, under different task requirements (and corresponding control specifications). This means that, to the extent that a task depends on control, it is not sufficient to consider the stimulus-response pair (*s, a*) alone. Rather, the task context *t* must be considered. We represent this as the triplet (a, *s, t*) and focus on the properties of *Q*(a | *s, t*) where, once again, *t* corresponds to the current task to be performed.

### B. Multitask Environments: Cross-Task Interactions

The formulation outlined above defines the informational demands of performing a single task. However, this becomes more complicated when the possibility of performing more than one task is introduced. In this section, we focus on the performance of multiple tasks simultaneously (i.e., “multitasking”)[123]. As an example, consider two tasks *t, t*^*′*^ respectively characterized by stimulus-response pairs (*s*_1_, a_1_, *t*_1_) and (*s*_2_, *a*_2_, *t*_2_). To address performance of these simultaneously, it would be natural to simply elaborate the formulation for performing each of them individually, given by Eq. 3, to include the joint stimulus and joint action pairs, along the lines of (*a*_1_, *a*_2_ | *s*_1_, *s*_2_) = − log_2_ *p*(*a*_1_, *a*_2_ | *s*_1_, *s*_2_, *t*_1_, *t*_2_). While this is a reasonable elaboration, it assumes the two tasks are independent of one another, without considering potential dependencies between stimuli, responses, and/or any internal representations that are shared by the two tasks. For example, in the case of the Stroop task, it does not take into account that color naming and word reading make use of the same phonological representations, precluding simultaneous execution of both tasks in response to an incongruent stimulus (i.e., it is not possible to accurately represent nor utter “red” and “green” at the same time).

As noted in the I, analyzing the interactions among more than just a few tasks in a neural network can become unmanageably complex. Thus, to address such interactions, we turn to the use of a simplified expression of the relationship among tasks in the form of a task graph (see Figure 1). The first simplification is to assume that all of the processing units in the neural network can be segmented into *sets*, each of which is represented as a node in the graph. For the moment we will focus on stimulus and response sets, designated S and R respectively, but the formalism generalizes to intermediate associative layers that we consider further on. In principle, a set can be an arbitrary collection of items (i.e., a stimulus set can be comprised of an arbitrary collection of stimuli, and similarly for responses). In practice, however, sets are assumed to correspond to naturally occurring dimensions along which stimulus features and response modalities are organized (e.g., colors, shapes, or locations; manual, verbal, or oculomotor responses). Following the definitions introduced in [57], we consider a task to be determined by a mapping *S* → *R*. This specification of a task generalizes the one used by Koechlin and Summerfield [13], by specifying dependencies among tasks in terms of the stimulus and/or response sets that they share. This, in turn, allows us to consider potential interactions among tasks at the level of the task graph.

As an example of how a task graph can be used to summarize and examine the interaction among a set of tasks fully defined at the neural network level, we consider the Stroop paradigm [64] referred to above, and shown as a task graph in Fig. 2a (corresponding to the format in Fig. 1b). In this simplified version, there are two sets of stimulus representations, one for colors, 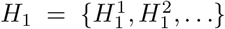 (e.g, 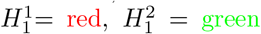, etc.) and one for words, 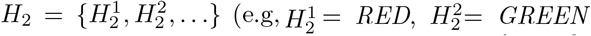. There is also a set of representations for verbal responses, 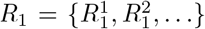 . In the color naming task, designated by the edge *f*_11_, participants are instructed to verbally report the color in which the stimulus is displayed. This is specified by the mapping: 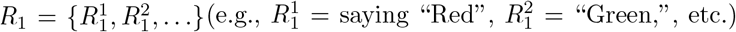 and 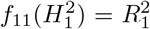. Similarly, the word reading task, in which participants are instructed to report the word while ignoring the color, and designated as *f*_11_, is specified by the mapping: 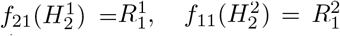. As noted above, participants are asked and, critically, are only *able* to successfully perform one of these tasks at a time; that is, either *f*_11_ or *f*_21_, due to their shared use of the response set. However, the task environment can be extended to permit multitasking by adding a third task, such as *word pointing*, in which participants are instructed to point in a specified direction in response to each word, as shown in Fig. 2b. Following [21], we refer to this as the *extended Stroop paradigm*. The word pointing task involves an additional response set for pointing in a given direction, 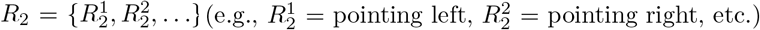, and the task is designated by the edge *f*_22_, that is specified by the mapping: 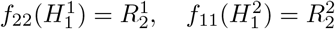. In the extended Stroop paradigm, participants can be instructed to perform any of the single tasks on their own (*f*_11_, *f*_21_, or *f*_22_), or to multitask color naming and word pointing (i.e., perform (*f*_11_) and f_22_) simultaneously). The latter is possible, at least in principle, because these two tasks do not share any nodes (i.e., they involve fully independent sets of stimuli and responses), unlike color naming and word reading (*f*_11_) and color naming (*f*_21_) which share a response set (*R*_1_). However, whether it can be done successfully *in practice* depends on the nature of *f*_21_. We will return to this point in Section III A.

**Figure 2.**
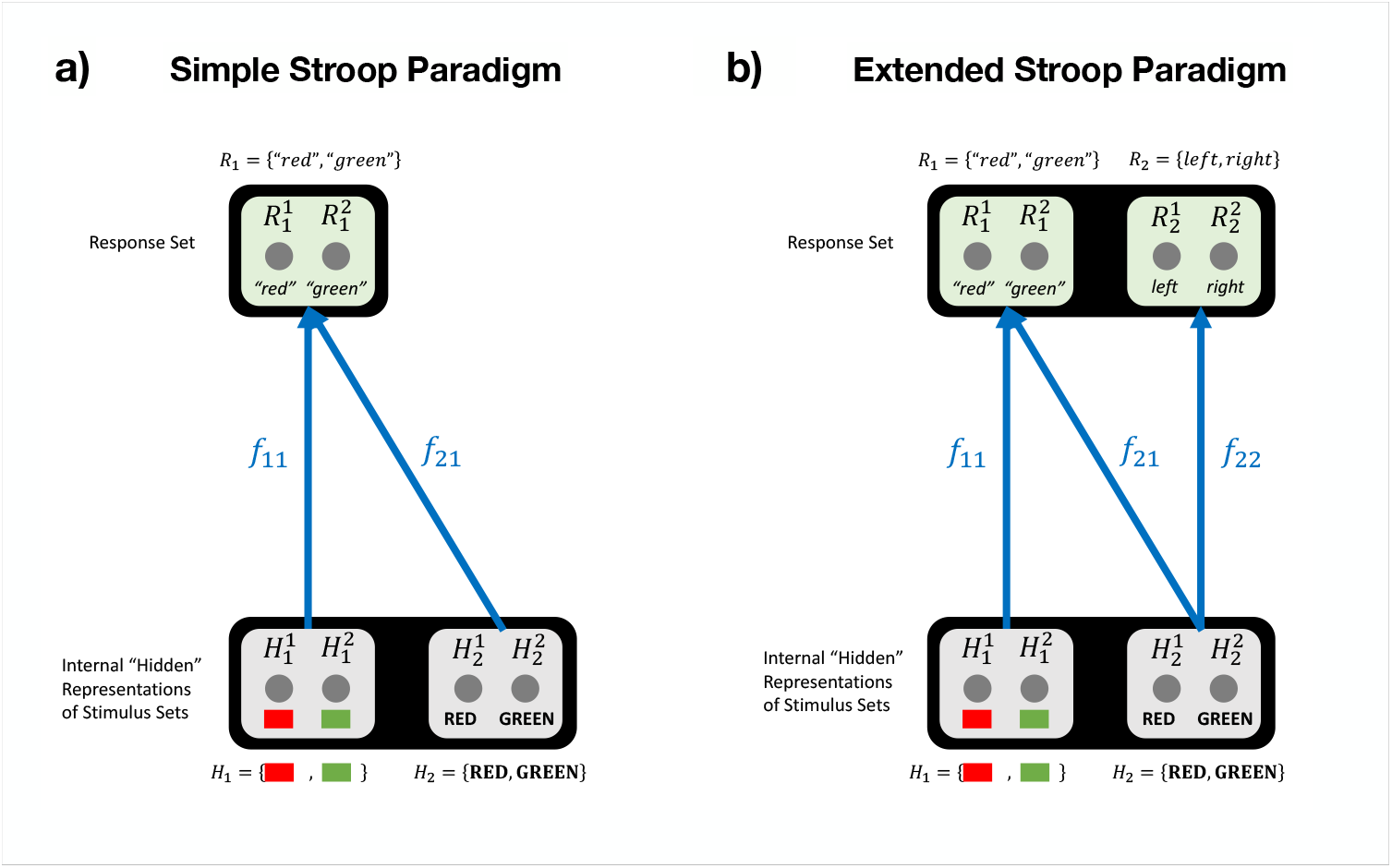
Tasks Graphs for the Stroop Paradigm. Examples of task graphs showing the relationship between representations of stimulus and response sets in: (a) simple version of the standard Stroop paradigm, involving color naming and word reading; and (b) an extended version that includes an additional word pointing task. Note that shaded boxes correspond to nodes of the task graph (light gray for input nodes and pale green for output nodes), with each node comprised of a set of processing units (dark gray circles) that represent individual stimuli or responses in the set, corresponding to the processing units of the neural network implementation, and edges representing the aggregate set of associations (connection weights) between processing units that comprise a task processing pathway (see text and Figure 1 for further explanation).

This example illustrates how the framework outlined above (and fully introduced in [57]) can be used to describe an architecture *U* = (𝒮, ℛ, 𝒯) capable of performing multiple tasks over a set of stimulus sets 𝒮 = {*S*_*i*_}_*i*∈{1,…*N* }_ and response sets ℛ = {*R*_*j*_ }_*j*∈{1,…*M* }_. Here, N and M are the number of distinct stimulus and response sets, respectively, and the tasks that can be performed are defined as a collection of mappings 𝒯 = { *f*_*ij*_ : *S*_*i*_ → *R*_*j*_ } between them (and in which any particular mapping f_*ij*_ might not exist for the pair (*i, j*)). This framework provides a formally rigorous characterization of tasks involving single, fixed mappings between stimuli and responses. While tasks of this form are a staple of cognitive research, as well as many applications in machine learning (such as classification tasks), they are obviously just a subset of the full range of tasks of which humans and machines are capable, that often involve complex *sequences* of processing. Below, and in the Discussion (Section V), we consider ways in which this framework can be extended to address more complex tasks, as well as initial efforts to do so. Nevertheless, even focusing on tasks involving single, fixed mappings, the framework can be used to formally analyze a number of factors that determine how interactions among tasks determine dependence on control, and that are likely to generalize to more complex tasks. Here, we explore three of these factors: (i) differences in the level of automaticity among tasks (determined by the relative strength of their processing pathways); (ii) the number of processing layers implementing the input-output mappings for the tasks (i.e., the depth of the network); and (iii) the potential for simultaneous parallel execution of multiple tasks (i.e., *concurrent multitasking*). In the section that follows, we consider how each of these factors determines the demands for control and the processing constraints with which this is associated.

### C. Functional relationship between neural network and task graph descriptions

As discussed above, and illustrated in Figure 2, the task structure identified by set 𝒯 can be represented as a task graph, with nodes defined by the sets *H*_*i*_ in one layer that encode representations of stimuli, and the sets {*R*_*i*_} in another layer that represents responses, linked by edges representing the mapping between them as defined by the task(s). This can be used to describe a neural network structure that can perform different tasks, by defining the strength of their pathways (represented as edges) required to perform individual tasks, and their relationship to one another. For tractability, previous work using this approach (e.g., [16, 19]) has treated edges (i.e., representing the sets of connections that associate stimulus representation sets and response sets) as binary, encoded in an adjacency matrix A in which the entry *a*_*ij*_ = 1 ⇔ ∃ *f*_*ij*_. This assumes that all tasks (and all stimulus-response associations within a task) are of equal strength (i.e., identical degree of automaticity). Here, we extend this approach to include weighted edges, used to represent the strength of associations between *H*_*i*_ and *R*_*j*_, and corresponding to the automaticity of each task, by assigning a weight ω_*ij*_ to a given stimulus-response set pair if an association exists between them (∈ ℝ ^+^ ⇔ ∃*f*_*ij*_), and 0 otherwise.[124]

#### Automatic efficacy and cost for single tasks

Using these definitions, we can formally express the effectiveness with which a task can be performed in the absence of control. We refer to this as *automatic efficacy* that, for a given task *f*_*ij*_ is defined as:

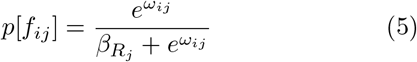

where ω_*ij*_ designates its strength (i.e., of the associations implementing the mappings that define the task), and β_*j*_ designates its baseline inhibition (i.e., determines the likelihood that a response unit *R*_*j*_ will exceed its threshold, and the corresponding action is taken). Note that, insofar as we are not interested in reflexes, we assume that by default the value of inhibition for all nodes is 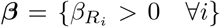 and typically much larger than the ω_*ij*_. That is, as discussed above, even strongly automatic processes (other than reflexes) require *some* amount of control to overcome baseline inhibition and elicit an overt response. Nevertheless, such processes can still have an automatic influence on processing in the absence of control. For example, this captures the influence of words on color naming that produces the Stroop effect as described above: word processing (labelled as f_11_ in Fig. 2a) is associated with an automatic efficacy (weight *w*_21_ >> *w*_11_) that is sufficient to allow orthographic stimuli to influence verbal responding even when no control is allocated to the word reading process, thus interfering with color naming (*f*_21_ in Fig. 2a).

It is important to note that performance efficacy as defined above should be considered from an information-theoretical perspective as the fraction of information available in support of performing a task relative to the amount that would ensure perfect performance. This should not be confused with the probability of actually responding correctly (i.e., accuracy), inasmuch as it does not take account of dynamical factors, such as the *integration* of information over time that are known to underlie performance in natural systems (e.g., see Note V D and Appendix A). Rather, it can be thought of as more closely related to signal strength (such as the drift rate in dynamical models [37, 38]), which contributes to but does not uniquely determine either accuracy or response time.

We also note that processing efficacy can be related directly to Equation 1 (following [13]) as:

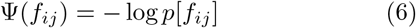

which formulates it as the cost associated with automatic performance. We use this formulation further below, together with a similar one for the cost associated with control-dependent processing, to consider both the overall cost of processing in various network configurations (Section II D) and how this can be optimized to maximize reward rate (Sections III and IV). *Task pathways in multilayer networks*. Finally, before considering the influence of control, we generalize the notion of automatic efficacy, and the corresponding cost, to cases in which tasks rely on pathways that span multiple layers of processing units in the network, as well as task environments in which multiple tasks are possible. Multilayer networks encompass both “deep” feed-forward networks that are widely used to address the complex mappings involved in naturalistic tasks such as computer vision ([25]), as well as recurrent networks used to address tasks that involve temporally extended and/or sequential processes [66–69]. For the purpose of illustration, we focus here on feed-forward networks, but the formalism can readily be extended to recurrent networks.

Consider a network architecture described by a multi-partite graph with L layers constituted by sets 𝒱_*l*_, with *l* = 0, …, *L* − 1. This can be transformed into a feed-forward network by allowing edges only between sets 𝒱^*l*^ and 𝒱^*l*+1^, in which each of these edge sets is called ε ^*l*^. It is natural to think each edge 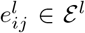 as a *sub-mapping* 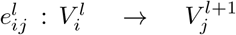 propagating inputs from internal set 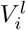 to internal set 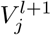. This can be thought as of a stack of bipartite networks (akin to those considered above) layered on top of one another, with the only condition that the first layer is *V* ^0^ = 𝒮, corresponding to stimuli, and the last is *V* ^*L*−1^ = ℛ, corresponding to the responses.

Accordingly, and following similar treatments in previous work (e.g., [32]), the definition of a task introduced above (in Section II B) can be generalized to a composition of sub-mappings from an input set to an output set along a specific path 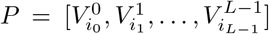 through the network. For example, the task of reading a word can be considered as a series of sub-mappings, from the visual input to orthographic representations of letters, from those to lexical representations of words [70, 71] and/or directly to corresponding phonological representations used to generate a verbal response (as in Figure 2). Such task pathways can be expressed formally as:

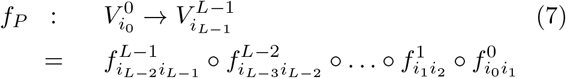

If we denote the set of all such tasks that can be performed by a given network as 𝒫, then the set of the processing pathways in the multitask network for performing those tasks can be written as: 𝒰 = ({𝒱^*l*^}, { ε ^*l*^}, 𝒫).

Using this framework, we can define the processing of task P to be successful if, at every step l along its pathway, the corresponding edge 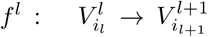 is successful in producing the associated response on unit 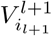. (Note that the the bipartite case discussed above is a simplified case of this more general definition, in which there are only stimulus representation and response sets and no internal layers). More specifically, the automatic efficacy for the task pathway *P* is given by:

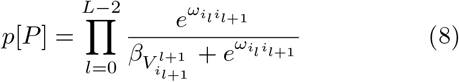

It is easy to see that, on the assumption that there is some finite amount of inhibition [125] associated with processing at each layer (represented by the β parameters) then, all other factors held constant, efficacy decreases with network depth due to the accumulated effects of β across layers as pathway length increases. This is illustrated in Figure 3a), which shows the efficacy p[P] for a single task path P processed through a network of growing depth for various possible β values (for simplicity, in this case, we set the β to the same value throughout a given network). As the depth increases, the probability decreases rapidly, implying that automatic processing of even a single task incurs greater reductions in processing efficacy in deeper networks. Consequently, the automatic processing cost Ψ(*P*) increases with depth (Fig. 3b, dashed lines) and inhibition (higher values of β). As we demonstrate in Section III B, increases in network depth are also associated with a greater likelihood of two tasks interfering with one another, yielding greater performance costs.

**Figure 3.**
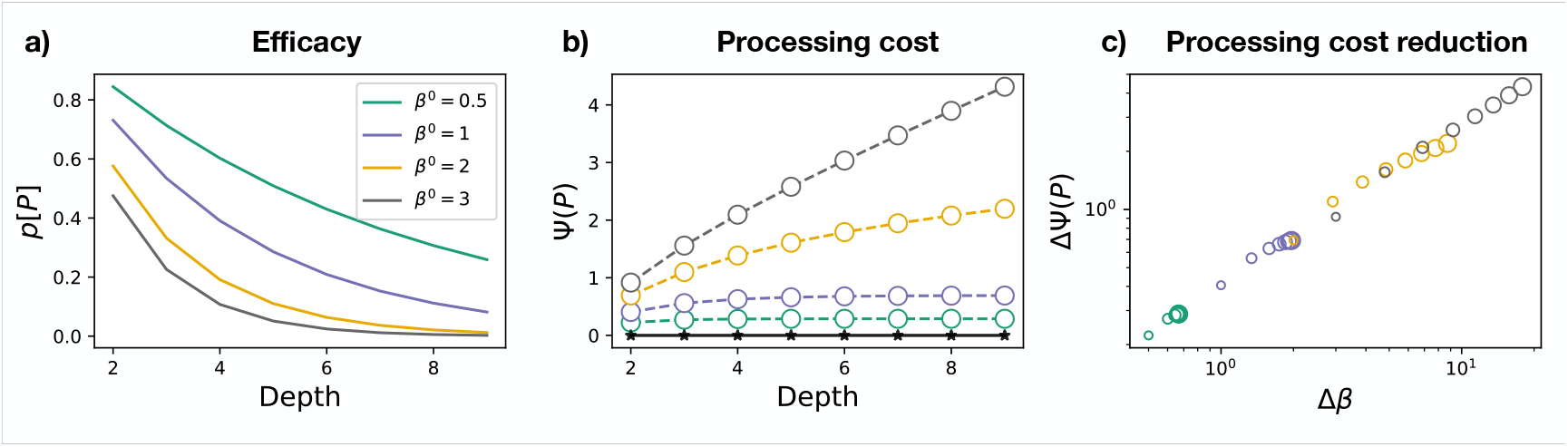
Performance cost and control for single tasks. (a) Efficacy *p*[*P*] for single task execution as a function of network depth for various values of *β*. (b) Automatic processing cost Ψ (solid lines) and ΔΨ (dots) for single task execution as a function of network depth for various values of *β*. Note that, for single tasks and sufficiently small values of *β*, the processing cost approaches and even reaches 0 (ΔΨ = Ψ). (c) Relationship between the processing cost reduction ΔΨ and the total amount of applied inhibition **Δ*β***, which represent the control applied to modulate the basal inibitions ***β***.

*Example: The Stroop Task Paradigm*

As an example of how the information-theoretic framework introduced above can be used to summarize task performance in a given network architecture, we compare this to the results of a neural network model that has been used to simulate the patterns of human performance in the Stroop paradigm [27]. Figure 4a) shows the two processing pathways in that model used to simulate the color naming and word reading tasks. Each pathway is comprised of a set of input units used to represent each of the two stimuli in each of the two stimulus dimensions (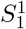 and 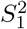 for the colors red and green; and 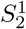 and 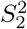 for the words RED and GREEN). The input representation units in each dimension project to a corresponding set of internal (hidden) units 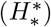 that, in turn, project, to units in the output layer of the network used to represent each of two verbal responses (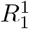 and 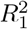, corresponding to “red” and “green” respectively). Thus, the network is comprised of two processing pathways, one for color stimuli and one for word stimuli, that converge on a common set of output units. The connections between processing units in Figure 4a represent the associations between each stimulus and each response that constitute the mappings defined by the task, with the sign (shown in black for positive and orange for negative) and weight (thickness) of the connection designating the strength of the association between individual stimulus and response pairs (i.e., their automaticity). Note that each stimulus is positively associated with a single response, and negatively with all others. This is consistent with a one-to-one mapping of stimuli to responses. The weight of the connections in the word reading pathway is greater than in the color naming pathway, capturing the greater automaticity of word reading relative to color naming. This network corresponds to the task graph shown in Figure 2a [126]

**Figure 4.**
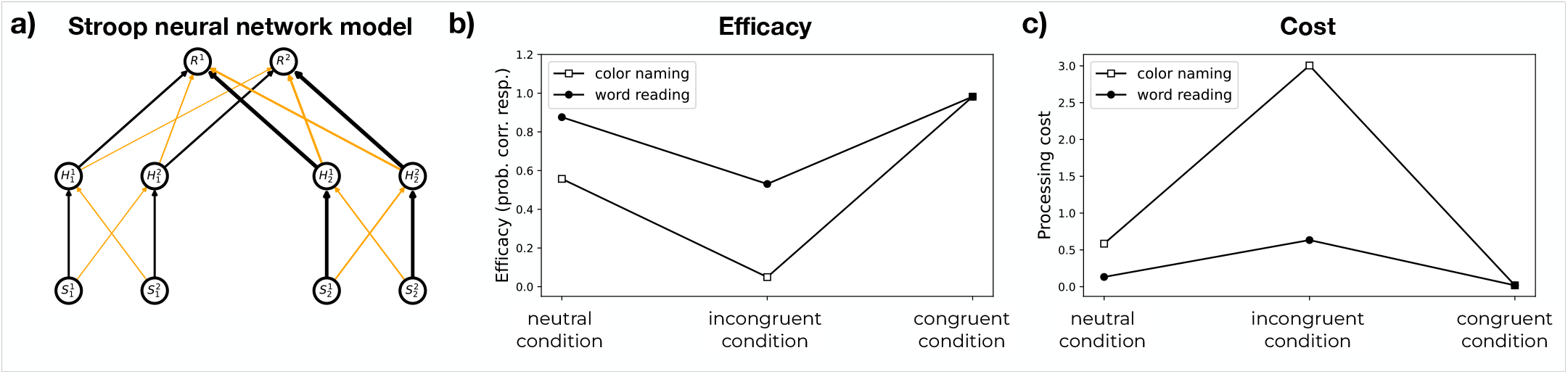
Information theoretic analysis of automatic performance in the Stroop paradigm. Panel a): Neural network model of automatic processing in the Stroop task (adapted from [27], showing the processing pathways for each task, but not the mechanism responsible for control). Excitatory connections are shown as black lines, inhibitory connections as orange lines, and line thickness represents connection weight (corresponding to |*ω*| at the network level; see Figure 1). The pathway on the left 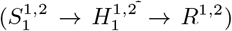 implements the stimulus-response mappings for the color naming task, and the pathway on the right 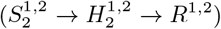 the mappings for word reading (see text for fuller description). All units have an inhibitory bias (corresponding to *β*) = 500. Panels b) and c): Efficacy values and processing costs, respectively, calculated for automatic performance under each of the three task conditions (see text) using the information theoretic formulation of the task corresponding to the network shown in Panel a) (see text and Equations 9-13). Network parameters are reported in Tables I in Appendix C.

This network can be used to simulate performance of the two tasks, color naming and word reading, each under the three possible stimulus conditions: *neutral*, in which relevant information is presented in only one dimension (e.g., for color naming, X’s shown in red [XXX] and, for word reading, the word red shown in black [**RED**]); *incongruent*, in which the color and word information disagree (e.g., GREEN); and *congruent*, in which they agree (e.g., RED). Simulations using this network produce results that qualitatively match those of human performance [27]. That is, they are characterized by four distinctive and robust effects: i) performance is better overall for word reading in all conditions; ii) word reading performance varies minimally across conditions, showing little effect if any of the color on the ability to respond to the word; iii) color naming is heavily affected by the word, with substantial interference in the incongruent condition and facilitation in the congruent condition, relative to control; iv) the interference effect is substantially greater in magnitude than the facilitation effect. The first three of these are consistent with the contention that word reading is more automatic (i.e., has a stronger processing pathway) than color naming, while the asymmetry of interference and facilitation effects has been attributed to nonlinearities in processing [27].

The original neural network model was designed to simulate both the *dynamics* (e.g. responses times) and *outcome* (i.e., response accuracy) of processing. The formalisms introduced above can be used to compute an information theoretic characterization of performance that, while not taking account of the dynamics (see Note V D), nevertheless accurately captures the effects of relative differences in the strength of the two processing pathways, and their interaction, on the outcome of performance. This can be done by first using (Equations 5-8) to compute the probability, for a given input, that only the correct response unit (and not the incorrect one) is activated. For example, in the neutral condition for color naming, if the stimulus is the color red, then 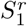 is the only input unit activated, and the desired propagation of activation (designated by →) is 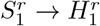 and 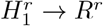, and *R*^*g*^ should *not* be activated (i.e., 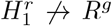). The probabilities for the corresponding activations under this condition are given by:

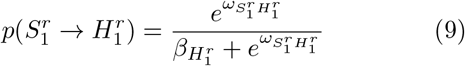

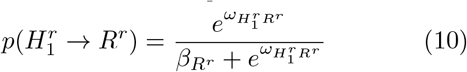

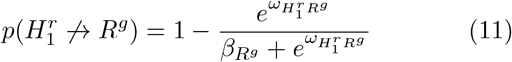

This can be used, in turn, to compute the automatic efficacy of performance for that input:

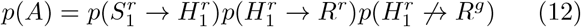

A similar computation can be performed for all of the other stimuli (i.e., in each of the three conditions for each of the two tasks). Note that, in the neutral condition used in the example above, only one source of input to the response units had to be computed. However, in the incongruent and congruent conditions, there are two sources of input. This can be quantified by computing the probability of the responses given all incoming connections. For example, in the incongruent condition, if the stimulus is comprised of 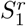 and 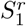, then the probability of responding to the color rather than the word (i.e., activating *R*^*r*^ rather than *R*^*g*^) is computed as:

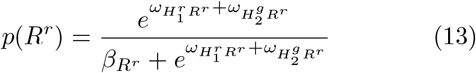

More generally, by calculating the activation probabilities for each unit given a particular input, and summing over all possible input patterns for a given task under each condition, we can determine the *automatic* processing efficacy and the processing costs for a given task under each condition. [127]. The results of these calculations are shown in Figure 4(b and c), assuming that the “correct” response to each stimulus is the one associated with the input in the task-relevant stimulus dimensions. Like the neural network model, the information-theoretic analysis closely aligns with the four canonical empirical results: i) word reading shows greater overall efficacy than color naming (compare the neutral condition for the two tasks); ii) word reading is only minimally impacted by information about the color of the stimulus (cf. word reading incongruent condition); iii) by contrast, color naming *is* profoundly impacted by information about the word (cf. color naming incongruent condition); and iv) interference is substantially greater than facilitation (compare the congruent and incongruent conditions against the neutral condition for each task).

Note, however, that this analysis deviates from the canonical effects associated with the Stroop paradigm in two ways. First, in the congruent condition, automatic efficacy is identical for color naming and word reading. On the surface, this is in contrast with the idea that word reading has greater efficacy than color naming; however, thus far, the information-theoretic analysis has only been applied to *automatic* efficacy, and has not taken into account any effects of control. Thus, for *both* tasks, performance is governed equally by all available information (i.e., *both* the color *and* word dimensions), and thus determined by the strongest pathway; in this case, that is word reading, and thus performance in the congruent condition is at “ceiling” irrespective of task. For similar reasons, the analysis indicates that the efficacy for color naming in the incongruent condition is at floor; that is, the network is unable to respond correctly, instead producing the word as the response. This result is consistent with the intuition (and easily verified prediction) that if an individual who has no familiarity with the Stroop paradigm is presented with an incongruent stimulus and asked to respond out loud *without any additional instructions* (i.e., no incentive to allocate control to process one dimension over the other), they are almost certain to respond to the word rather the color. In the Stroop model [27], both the ability to name the color, performance differences between color naming and word reading in the congruent condition, are explained by the allocation of control, to which we turn in the next section.

In summary, we have shown that our formalism, applied to task processing described at the neural network level, is able to reproduce characteristic features of performance in the Stroop paradigm. However, as noted at the outset, analysis at the neural network level – that takes account of both the full details of stimulus-response associations across all tasks as well as the dynamics of processing – can quickly become intractable when considering potential interactions among multiple tasks, each of which may involve a multiplicity of stimuli and responses. Accordingly, to examine interactions among tasks, and how this impacts the requirements for control and multi-tasking capabilities, we turn to an analysis at the level of the task graph. We use this to consider task interactions in terms of the interference produced by patterns of task “dependencies” (that we define in Section III A 1 below). First, however, we develop the construct of control at this level of analysis.

### D. Modulation and Control Efficacy

To address the influence of control at the task graph level, we add we add a parameter **∆*β*** to the expression for efficacy in Eq. 5, that is a vector with a value for each node of the task graph that modifies its corresponding baseline ***β***. Thus, for a given node j receiving an input from a set of nodes I, the activation probability can be written as:

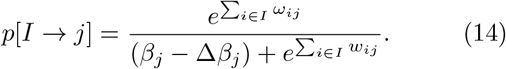

The role of the ∆β_*j*_ is to diminish the inhibition of j, hence facilitating j’s activation. Although **∆*β*** is implemented as an additive parameter, its interaction with the non-linearity of the activation function has a multiplicative, gating-like effect on processing: since baseline β is assumed to be high, processing units associated with the units (e.g., see Figure 2) are not only inhibited, but also in an insensitive range of their activation function (i.e., changes to other inputs have little effect on the unit); by diminishing β, **∆*β*** places units in a more sensitive range of their activation function, and thus renders them more responsive to other inputs (see [27] for a more detailed discussion). By modulating the sensitivity of processing units in a task pathway, the **∆*β*** can be used to control the efficacy of that task, in a manner similar to the multiplicative (“gating”) effects used to implement control in models with linear processing units (e.g., [75, 76]). For example, where the activation function is a sigmoid (e.g., logistic or tanh) and the net input and β appear in the exponent, **∆*β*** has the effect of adding to the exponent, which is comparable to multiplying the net input when the activation function is linear.

To the extent that each ∆β_*j*_ applies to a unit in the network, then its value applies to all processing units represented by that node (see Figures 2 and 1). This aligns with the simplifying assumption that processing units are arranged into sets that represent similar information, and that control operates uniformly over such sets of units (i.e., over a node), and independently of others. This approach has been used productively to model the effects of control in tasks that align with well-defined and dissociable categories of information, such as colors and words in the Stroop task [21, 27, 29, 77, 78]. More generally, however, the representations required to perform different tasks may not be so discretely separable. Nevertheless, the ability to perform different tasks based on similar information requires that there be *some* basis on which distinctions can be made, usually at a higher level of abstraction, that can be exploited by the control system to perform the different tasks (see [79] for recent work that has begun to address how the simplifying representational assumptions made here can be generalized to address control over more graded distinctions among distributed representation reflecting more complex forms of semantic structure. Figure 4b and c (solid lines) show the automatic efficacy of performance for the color naming task (see Appendix for the specific **∆*β*** parameters), which aligns qualitatively with the effects in a neural network model of the Stroop task [27], including the observation that the magnitude of interference is influenced by the relative strength of one task with respect to the other (e.g., word reading interferes more with color naming than the converse and, accordingly, color naming cannot be performed for incongruent stimuli without control).

For the case of multiple tasks, the efficacy for a given task pathway *P* (again with node *j* = *i*+1) can be written as:

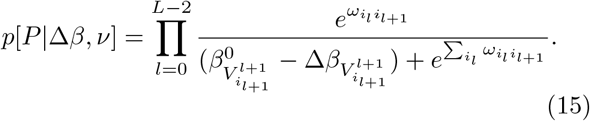

The **∆*β*** parameters in this expression implement the function of control discussed above. That is, ∆β_*i*_ for node i mediates its effects by counteracting the intrinsic inhibition imposed by the corresponding β_*i*_, thus disinhibiting unit i. This not only makes it more likely for weaker pathways, which are more influenced by intrinsic inhibition, to reach their threshold for responding, but also allows them to compete more effectively with stronger pathways (i.e., ones with greater automatic efficacy) to which control has not been allocated (but that are less impacted by their own intrinsic inhibition). Since the **∆*β*** parameters account for changes in efficacy due to the allocation of control, we refer to their contribution to the overall efficacy p as the *controlled efficacy* component.

Note that, in addition to **∆*β***, the expression for pathway efficacy (Eq. 15) is dependent on another parameter, *v*. This represents another form of control that has its primary effect on the interaction between tasks in multitask networks, and that we consider in Section III B. In single task networks, however, *v* has effects qualitatively similar to those of **∆*β***, and thus we ignore it for the remainder of this section.

In general, **∆*β*** can be thought of as reflecting the influence of task representations used for control—that themselves are activity-based—and thus subject to rapid updating. Here, we do not directly consider the mechanisms responsible for the activating and updating (nor the learning) of such representations. However, we assume that, as suggested by other work, they are subject to the same principles of learning, representation, and processing as the ones discussed here, simply applied at a higher (e.g., more abstract) level of representation [12, 27, 79, 80]. We return to this point further on, where we discuss the *acquisition* of automaticity. First, however, we focus on the effects that the elements of **∆*β*** have on overall efficacy p, by treating a given **∆*β*** as a *control policy* and considering how it can be adjusted to optimize processing.

The simplest, but the most important effect is the interaction of **∆*β*** with the β and ω parameters that determine pathway strength (i.e., automatic efficacy), allowing weaker pathways to compete more effectively with stronger ones: the greater the disparity of automatic parameters between two pathways in competition, the more the weaker one relies on its ∆β_*i*_’s for execution.

Using Eq. 15, we can formulate the information cost Ψ for P associated with a control policy **∆*β*** as:

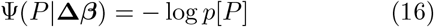

and the cost reduction ∆Ψ induced by that control policy as:

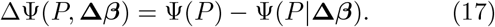

The control policy **∆*β*** can then be optimized, by finding elements that minimize the processing cost Ψ (and thereby maximize processing efficacy p) for a given task pathway P. Figure 3b shows that for single tasks, it is always possible to find the set of control values **∆*β***^∗^ for the corresponding pathway that bring the controlled processing cost to zero, Ψ^∗^ = 0 (Fig. 3b, black stars), which in turn implies ∆Ψ = Ψ −Ψ^∗^ = Ψ (Fig. 3b, coloured dots). This is trivially done by setting ∆β_*j*_ = β_*j*_, which in turn makes all efficiencies 1 and thus all costs 0. However, this requires progressively more control to be applied as the network becomes deeper, which implies that the requirement for control **∆*β***^**∗**^ grows with network depth, proportionally to the growth in processing costs (Fig. 3c). Furthermore, in the next section we will see that, even with an unlimited budget for control, it is not always possible to bring performance costs to zero when *multiple* tasks must be performed. Moreover, as control engages more task pathways, there is a greater potential for those task pathways to interfere.

## III. OPTIMIZATION OF CONTROL AT THE TASK GRAPH LEVEL IN MULTITASK ENVIRONMENTS

In the previous section we affirmed that the treatment of automatic and controlled efficacy in information-theoretic terms aligns closely with previous work using neural networks to model the effects of pathway strength (automaticity) and activation-based modulation (control) under conditions in which a *single* task must be performed, but its pathway shares a set of representations with another task pathway that may have greater or less automatic efficacy (e.g., the output representations shared by color naming and word reading in the Stroop paradigm). Here, we broaden the scope of the approach to address the optimization of control in situations where more than one task must be performed. To do so, we introduce some additional simplifications.

Following similar efforts [14, 16, 19, 21, 32, 57], the first simplification is to assume that for tasks that share representations, the inputs relevant to each task are always associated with different specific representations in the shared set (as in the incongruent condition of the Stroop task; see Figure 4b). This is motivated by two considerations. First, as noted above, the costs associated with incongruence (interference) are generally much greater than the benefits associated with congruence (facilitation; e.g., see Figure 4, Panels b and c), and therefore likely have a greater impact on processing. Furthermore, in realistic environments, incongruence among independent stimulus dimensions is much more likely than congruence [128]. For both of these reasons, optimization of control in multitask settings is likely to be driven more strongly by the interfering effects of incongruent stimuli than the facilitative effects of congruent ones.

The second simplification follows the approach taken to the formulations of efficacy above (e.g., Equations 12 and 15), by focusing on effects at the graph level through the construction of a *task graph* (along the lines used to depict the Stroop paradigm in Figure 2). In this graph, each node corresponds to a *set of processing units* used to represent similar information (as discussed in Section II C), and edges correspond to the *mappings among them* that define each task. For example, the two hidden units per stimulus dimension (colors and words) in the model of the Stroop paradigm shown in Figure 4a (and corresponding to the model shown in the left of Figure 1) are each summarized by a node, *H*_1_ and *H*_2_ respectively (shown on the right in Figure 1). The shared set of verbal responses units are represented by *R*_1_, and the strengths of the mappings in each pathway by the edges (*f*_11_ and *f*_12_, respectively [129]. Note that the state of each node in the task graph—rather than representing the activation of stimulus- or response-specific information—represents the likelihood that information corresponding to a particular type of information is processed.

### A. Automatic Multitask Processing

Thus far, we have considered processing efficacy when the network is asked to perform only a single task at a time. However, the formalism outlined above can be used to describe a network architecture 𝒰 capable of performing multiple tasks, and to analyze how the architecture impacts efficacy of task performance. As discussed in the Introduction, reward can sometimes be optimized if more than one task is performed at a time, by performing them in parallel; that is, through *concurrent multitasking*. For a given network architecture, parallel execution can substantially increase the reward rate for some combinations of tasks (that do not share representations). However, for other combinations (that *do* share representations), parallel execution may be disadvantageous by introducing the potential for interference, which can compromise performance and thus diminish reward rate. Such combinations demand serial execution and concomitant dependence on control. Here, we consider how the analysis of efficacy can be applied to the question of when (and for which tasks) performance should be parallelized, by formalizing the automatic efficacy of parallel execution (multitasking) for a given set of tasks, as a benchmark against which the value of serial processing and the demands for control can be compared (that we address in the next section). We start by considering single-layered networks, in which representational sharing is restricted to stimulus and/or response sets, and use this to define different forms of dependence that can arise among tasks as a consequence of different configurations of sharing. We then build on this formalization to examine more complex, multilayered networks.

#### 1. Single Layer Task Graphs

In Figure 5, we show the combinations of tasks that can be formed from a pair of stimulus and response sets, arranged from left to right according to the extent of sharing of sets between tasks. In the leftmost configuration (*a*), the two tasks do not share any stimulus or response sets, and thus are independent of one another and do not pose a risk of interference, allowing them to be performed in parallel. All of the other configurations exhibit various forms of dependence among tasks, that pose the risk of interference, though for different reasons and with different consequences. We first consider these qualitatively—in terms of a distinction between *structural* versus *functional* dependence—and then quantify them using the formalisms introduced above.

**Figure 5.**
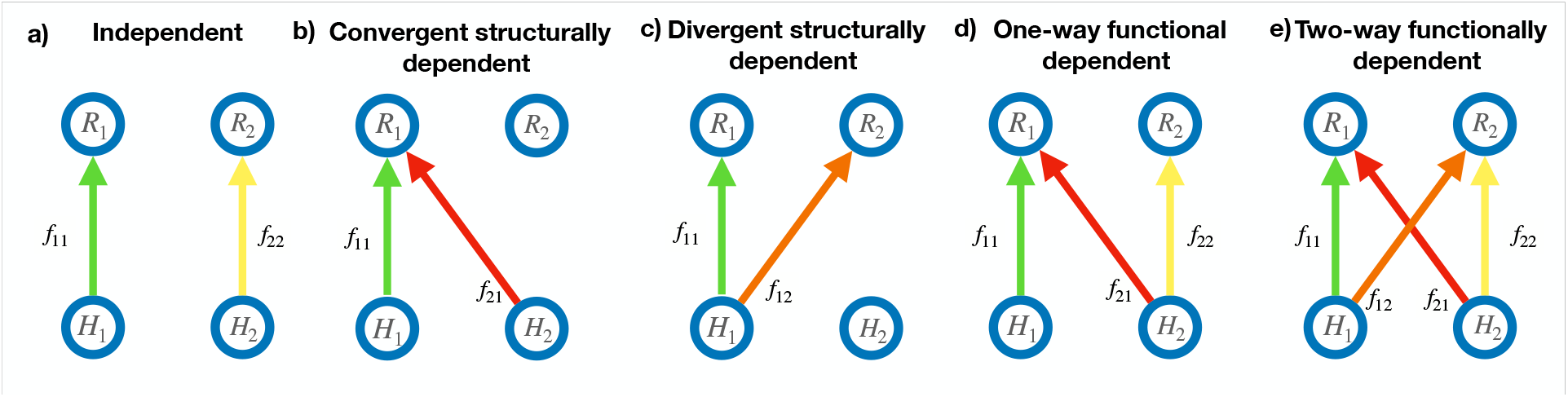
Possible configurations of interference among tasks. Blue nodes show stimulus and response sets, and colored edges show tasks formed by various mappings between them. (Note that here, as in Figure 1, we label the input nodes of the task graph as *H*_*i*_, on the assumption that each receives a mapping from a different stimulus set (as in the model of the Stroop paradigm shown in Figure 4), and thus is sufficient to capture the neural network-level effects of shared representations at the hidden and/or response levels.) Green and yellow edges designate tasks that do not share representations with (i.e., are independent of) one another and thus can be performed in parallel. Red edges designate mappings that share a set of representations with another task, and thus either introduce dependencies among those tasks (panels b, d and e), or do not constitute a legal (independent) task (panel c; see Section II B or [57]). Different panels show examples of mappings associated with different types of dependencies (see text), arranged from left to right according to extent of representational sharing.

##### Structural dependence

The problem of interference arises in the simplest form when a pair of tasks share the same response set (configuration *b* in Figure 5), as in the case of color naming and word reading tasks in the Stroop paradigm (see Fig. 2a and Figure 4a). This is because the inputs for the two tasks may make disparate demands of their shared response set (e.g., for the stimulus GREEN, say “red” in response to the color versus “green” in response to the word). This reflects two fundamental assumptions about the definition of tasks and the nature of the architecture (formalized in [57]): i) tasks are defined by the ability to sample a stimulus for processing *independently* of any other task; ii) the units in a given set (stimulus, internal or response) cannot simultaneously represent different information at the same time. Furthermore, as noted above (Section II B) and in [57]), independent sampling along stimulus dimensions means that the likelihood of an incongruent stimulus is much greater than of a congruent stimulus, and thus the risk of conflict is high for tasks that share a set of representations. We refer to such sharing of representations by two or more tasks, exemplified by the configuration shown in Panel b), as *structural dependence* [21, 83, 84], as it directly reflects a structural feature of the graph architecture.

As just suggested, structural dependence can also arise from a shared stimulus set if it maps to different response sets, as shown in Figure 5c. While this configuration would not seem to pose the risk of conflict (since it maps to independent response sets), it poses another, *conceptual*, problem. Distinct tasks are defined by the ability to sample task-relevant stimuli independently of one another, a requirement that cannot be met if two tasks share the same stimulus set. For example, imagine that one task is to name the color of the stimulus, and another is to point at a matching color patch in another part of the display. It is not possible to isolate these as separate tasks and perform them simultaneously, since it is not possible for the target stimulus to be two different colors at the same time (that is, to sample the stimuli for each task independently of the other). Conversely, if the *same* stimulus is always used for the two colors (i.e., the stimuli are not sampled independently for the two tasks), then what is labeled as two tasks could just as naturally be thought of as a *single* task that simply has a more complex response. For this reason, we treat two pathways that share a stimulus set as a form of structural dependence that, in this case, precludes considering their simultaneous execution as a form of genuine multitasking performance [130].

##### Functional dependence

A subtler case of dependence arises when two tasks do not share any stimulus or response sets (such the tasks shown in green and yellow in Figure 5d and e) or, more generally, even any internal representations—that is, they are not structurally dependent—but one shares a stimulus set and the other shares a response set with a third task (shown in red and orange). In this case, any attempt to simultaneously perform the fist two tasks inadvertently invokes the third task. That is, even though the first two tasks are not themselves structurally dependent, they are still at risk of the functional consequences of interference as a result of their mutual structural dependence on a third task. Accordingly, we refer to the first two tasks as being *functionally* dependent. The same applies for the sharing of internal representations (in place of or in addition to stimulus and/or response sets), so long as the same criteria apply: they are not shared directly by the first two tasks but, by disjoint sharing with each, they comprise a third task.

The consequences of functional dependence are illustrated by color naming and word pointing in the extended Stroop paradigm discussed above (Fig. 2b). To simultaneously perform the color naming and word pointing tasks (*f*_11_, *f*_22_ in Fig. 2b, respectively), control must be allocated to the color and word stimulus sets (e.g. by increasing 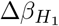 and 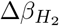, respectively), as well as to the verbal and manual response sets (by increasing 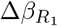 and 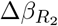). Note, however, that doing so allows information to flow from the stimulus set for words to the response set for verbal responses via *w*_21_, inadvertently engaging the word reading task. For incongruent stimuli, this will not just prolong responding, but will prevent an accurate response altogether. For example, in the standard Stroop task, when color naming is the only task being performed, information from a conflicting word may partially activate representation in the word stimulus set, which will compete with and thereby slow down processing of the verbal response to the color; however, since no control has been allocated to the word stimulus set, that information will not be active enough to determine the actual response. In contrast, when the word pointing task is performed simultaneously, and therefore control *is* allocated to the word stimulus set, it will now determine the response since its connection to the verbal response set is stronger than the one for colors (Fig. 4b). Thus, even though color naming and word pointing are not *structurally* dependent (they involve independent stimulus and response sets), they are *functionally* dependent by way of word reading. As a consequence, without a change in graph configuration (that we consider further on, in Section IV B), the graph cannot simultaneously and accurately perform (i.e., *multitask*) both color naming and word pointing. This is consistent with empirical findings that, without extensive practice, people are incapable of performing these two tasks simultaneously [21] (see [53] for another example). *Formal analysis of multitasking efficacy*. The risks of interference associated with multitasking performance, exemplified by the various types of dependence shown in Fig. 5, can be generalized to more complex bipartite graphs by defining a multitask combination as a set of tasks 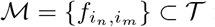 from an arbitrarily complex 𝒰 [57]. Simultaneous performance of the tasks in ℳ entails the activation of internal representations *H* _ℳ_ associated with the stimulus sets for all of those tasks, and the corresponding responses *R* _ℳ_ from the response sets for those tasks. Based on this, we can then identify the full set of tasks engaged by ℳ —that is, including any additional tasks inadvertently engaged due to functional dependence among those in *R* _ℳ_ —as the extended multitask 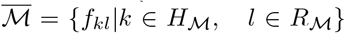, where by construction 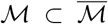. This is because, as discussed above, when the stimulus and response sets of ℳ are activated, all tasks that map representations *H* _ℳ_ to *R* _ℳ_ (i.e., those responsible for any functional dependence among them) are also engaged. For example, if multitasking is attempted for the green and yellow tasks in Figure 5d (i.e., ℳ = {*f*_11_, f_22_ }), then the red task will join them in comprising the extended multitask 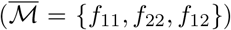. From the definition of 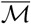, it follows that the probability of correctly performing any task *f*_*ij*_ ∈ ℳ depends on the extent to which the probability of its output response *R*_*j*_ depends on H_*i*_ relative to any other input to the graph H_*k*_ via the other tasks associated with 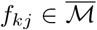 Accordingly, the automatic efficacy *p*[*f*_*ij*_| ℳ] of task *f*_*ij*_ in multitask *ℳ* can be expressed as:

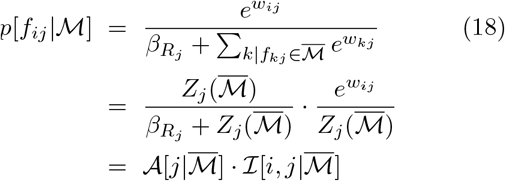

where 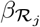 encodes the probability that the node *R*_*j*_ will not respond to any of the activated input set ℋ _ℳ_, and the term

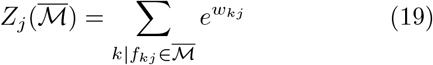

summarizes all edges originating from input nodes incident to node *j* in the context of the extended multitask set 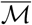. The function 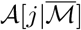 then corresponds to the probability that node *j* responds (i.e. is activated) in the context of multitask 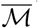, and 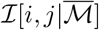 the probability that the input coming from the preceding node i is influenced by *j*.

Note that the difference between Eqs. 5 and 18 is the dependence of *Z*_*j*_ on 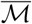, which in turn quantifies the effect of interference between shared representations among tasks.

Using Eq. 18, the performance cost of task *f*_*ij*_ in multitask ℳ is:

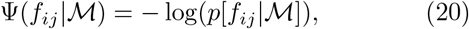

and thus the total performance cost of ℳ is:

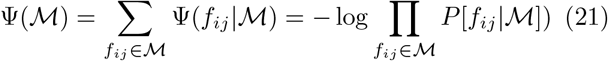

##### Examples

As an illustration of how Eq. 21 can be used to quantify the multitasking performance of different task configurations, we apply it to the ones shown in Figure 5 (we consider more complex examples in Appendix B). For the configuration shown in 5*a*, the two tasks are independent, and thus the multitask efficacy Ψ(ℳ) is simply the product of their individual efficacies, which can be computed using the expression for a single task (Eq. 14). However, for all of the other configurations, structural and/or functional dependence plays a role.

Configuration 5*b* shows a case in which two tasks (*f*_11_ and *f*_21_) are structurally dependent due their shared response set (*R*_1_). As a consequence, the individual efficacies of the two tasks,

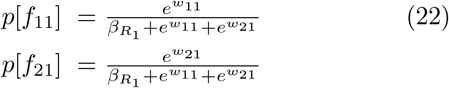

are in tension. If one is stronger (i.e., has a higher *ω*) than the other, then that one will prevail to an extent determined by the asymmetry, and only one of the two tasks can be performed reliably, as in the case of color naming and word reading in the standard Stroop paradigm.

In configuration 5*c*, the two tasks are structurally dependent due to a shared *stimulus* set. As discussed above, this makes it impossible to independently choose stimuli for the two tasks, and thus only one of the two tasks can be performed at a given time.

Configuration 5*d* and *e* illustrate cases in which two tasks (*f*_11_ and *f*_22_, shown in green and yellow, respectively) are *structurally in*dependent, but *functionally* dependent by way of either one (*f*_21_) or two (also *f*_21_) other tasks in the graph (shown in red). In these cases, even if a multitask is restricted to just tasks *f*_12_ and *f*_22_, their efficacies suffer by an amount dependent on the strength of the task(s) responsible for their *functional* dependence. This is illustrated in Figure 6. The top panel plots the efficacies for tasks *f*_12_ and *f*_22_ in the configuration shown in 5*d*, as a function of the strength (*ω*) of task *f*_12_ responsible for the functional dependence between them. Since *R*_2_ is not shared, the efficacy of *f*_22_ is simply:

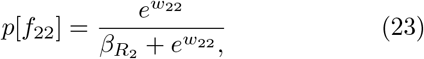

which is unaffected by *ω*_21_(flat yellow line in top panel of Figure 6). However, since *R*_1_ is shared between *f*_11_ and *f*_21_, they are structurally dependent, and their efficacies are the same as those for the corresponding tasks shown task in Figure 5*b*, given by Eq. 22 (and plotted, respectively, as the green and red lines in the top panel of Figure 6): As *ω*_21_ increases, the efficacy of *f*_21_ increases, and thus the efficacy of *f*_12_ decreases. A similar pattern is seen for the configuration shown in Figure 5*e*. However, in this case, since *R*_2_ is now shared between *f*_12_ and *f*_22_, they too are now structurally dependent, and thus the efficacy for *f*_22_ is now also compromised (as is the additional task *f*_12_ in this configuration). Note that, since neither is dependent on ω_21_, they are not affected by it (i.e., their plot is flat, with the height determined by *ω*_12_ = *ω*_22_ = 1 used in this example. These results provide a formal account of previous simulation results with non-linear neural networks, demonstrating that increases in the strength of interfering tasks (e.g., via training), can negatively impact multitasking performance for functionally dependent tasks [20]. Appendix B provides a more detailed characterization of the efficacies in these different configurations.

**Figure 6.**
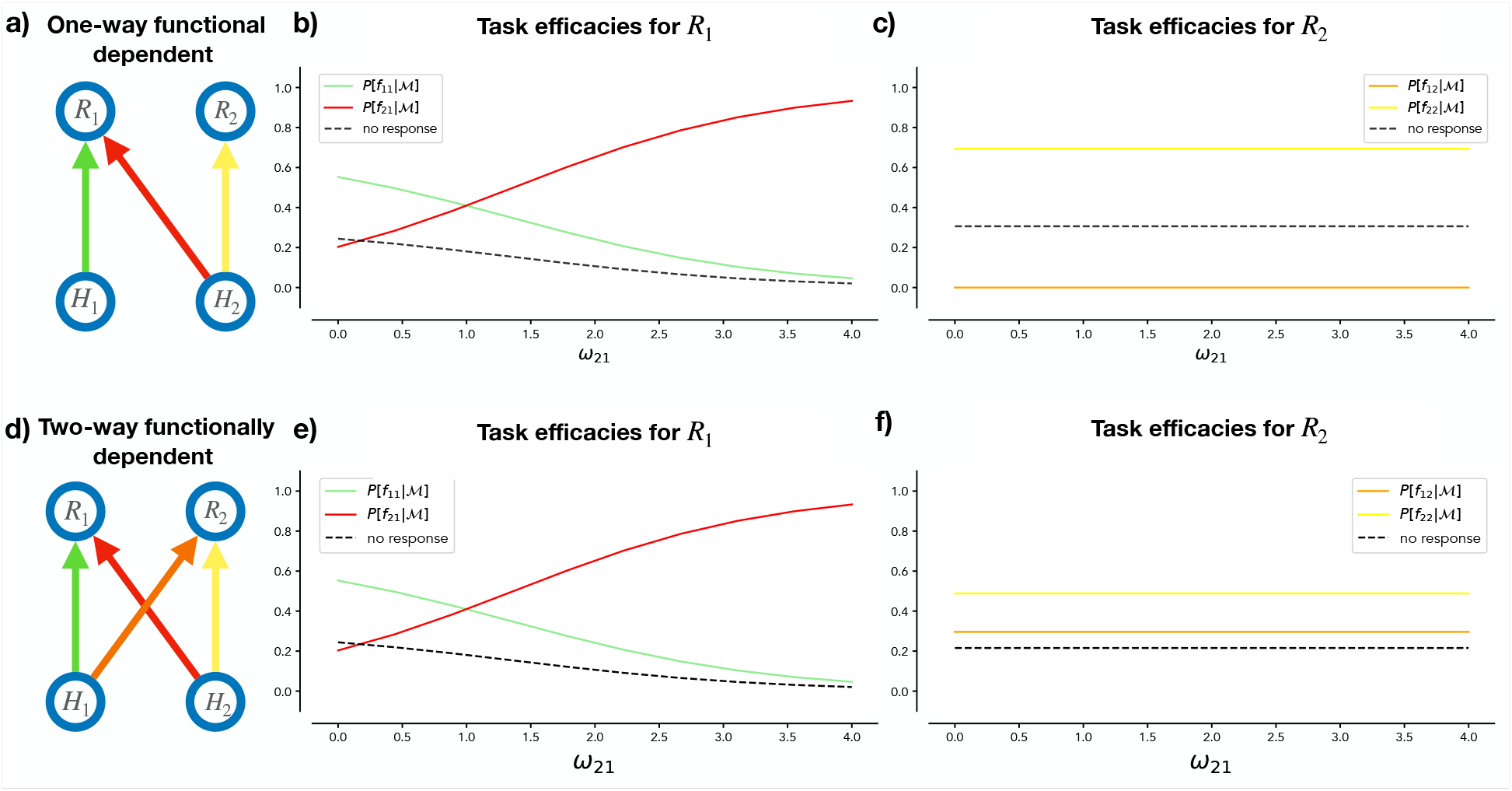
Task efficacies in multitask settings. Efficacies for individual tasks (calculated using Eq. 18) in the multitask settings shown in Figure 5*d* (reproduced in panel a) and 5*e* (reproduced in panel d), as a function of the strength (*ω*) of the tasks (shown in red) responsible for structural and/or functional interference in each configuration. Panels b) and c) refer to the configuration in panel a). Panels e) and f) refer to the configuration in panel d). The efficacy curves for each task are shown in the corresponding colors in the plots. For completeness and comparison with the bottom row, in the top row right panel we show also the curve for *f*_12_ although it is zero for all *ω*_21_ values. Note that in this case the presence of *f*_12_ changes the efficacy for *f*_22_. For these examples, *β* = 1.2, *ω*_11_ = *ω*_22_ = 1, *ω*_12_ = .5.

#### 2. Multilayer Task Graphs

The three kinds of relationships between tasks— independence, structural dependence, and functional dependence—introduced above with respect to two-layered task graphs, can be extended to multilayered graphs (i.e., from bipartite to multipartite graphs), as shown in Figure 7. A pair of tasks is considered independent if their pathways do not share any nodes, nor are they linked by nodes of any other pathways in the graph (as shown by the green and orange paths in 7*a*); tasks are structurally dependent if they share any nodes (Fig. 7*b*); and they are functionally dependent if they do not share any nodes directly (Fig. 7*c*), but are linked by links of any other pathway in the graph.

**Figure 7.**
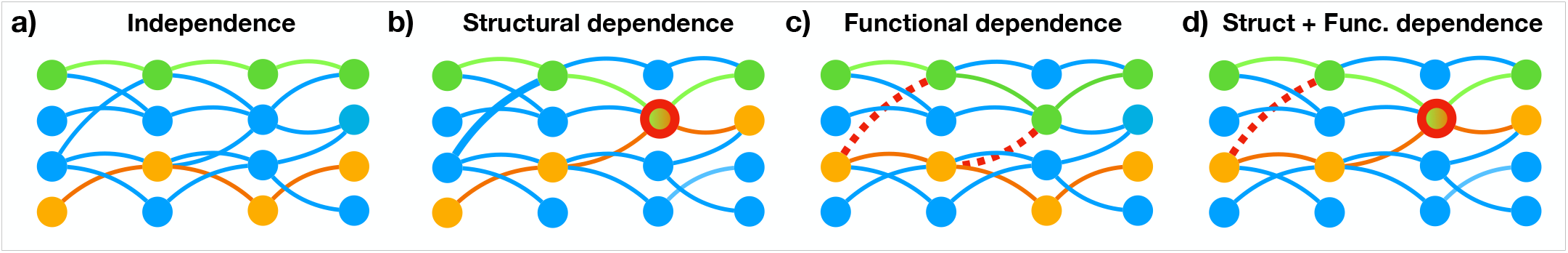
Dependencies among task pathways in multilayered task graphs. Examples of two task pathways (green and orange) in a task graph that also implements other task pathways (blue). Panels show configurations illustrating different dependency relationships between the green and orange pathways: a) independence; b) structural dependence, due to a shared node (outlined in red); c) functional dependence, due to other pathways that connect them (red dashed edges); and d) both types of dependence.

Note that, in the multipartite case, the dependence between a pair of tasks may not be limited to just structural or functional (e.g., 7*b* and 7*c*), but may involve both forms of dependence as shown in 7*d*. In general, the opportunities for dependence between tasks—and the attendant risks of interference—are considerably greater in a multilayered graph, that has been analyzed in previous work ([19, 32]). That work has similarly exploited the representation of neural networks in the form of task graphs, analyzed the consequences of structural and functional interference in terms of the matching and induced matching problems, and identified the maximum independent set as a way of determining the maximum number of independent tasks supported by the neural network. In particular, such work has shown that this number grows dramatically sublinearly with network size [14, 16], including depth [19, 32]. Our analysis reaffirms this observation, further suggesting that the effect of network depth is due largely to the increase in structural dependence (see Figure 8. Nevertheless, not surprisingly, functional dependence also remains a problem with deeper networks. Critically, and consistent with previous results, the combined effects of structural and functional dependence quickly become overwhelmingly prevalent, even for relatively shallow graphs (3-4 layers). This implies that, with an increase in the size of the graphs, the capacity for multitasking diminishes as the likelihood for cross-task interference grows, underscoring the importance of control to which we turn next.

**Figure 8.**
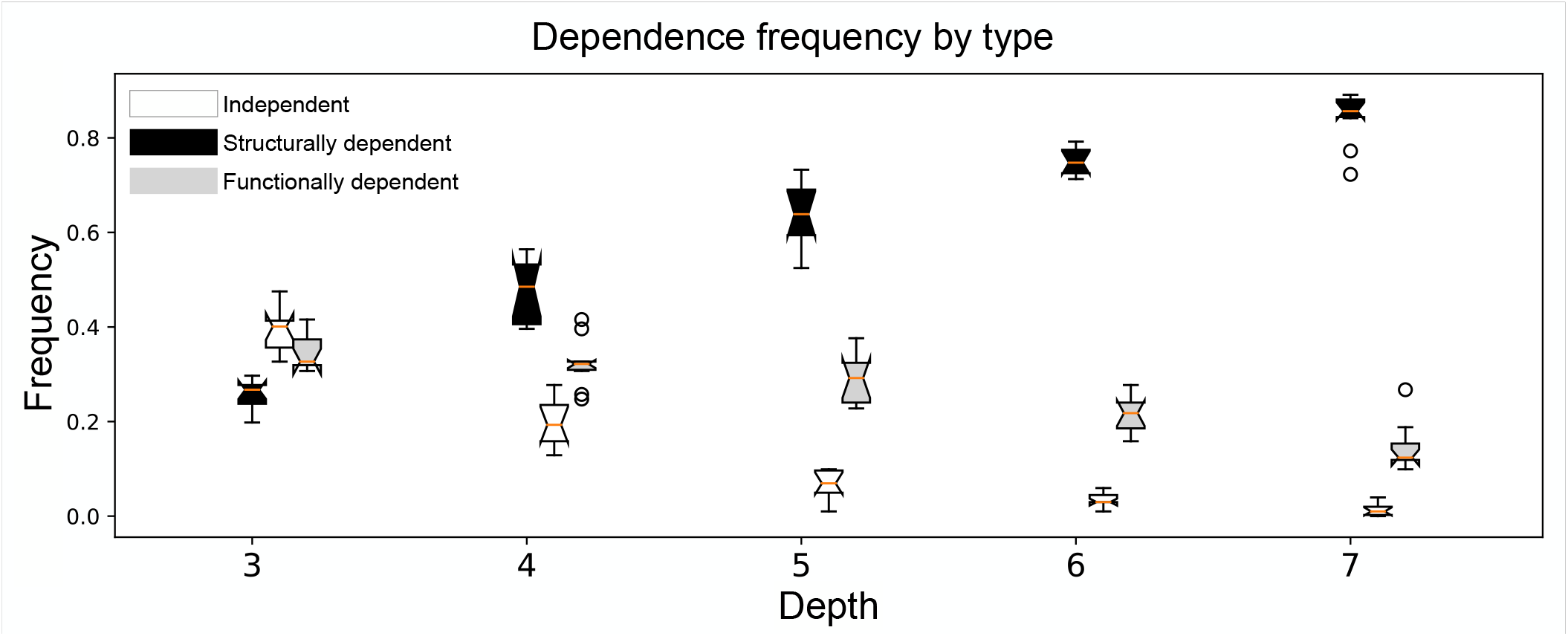
Frequency of dependency types for multitasks as a function of graph depth. Each point corresponds to the mean proportion (and its standard deviation) of multitasks composed by two tasks that are independent (white), structurally dependent (black), and functionally dependent (gray). Results are for 1000 simple graphs with 10 nodes per layer and edges between units across adjacent layers assigned uniformly at random, with a fixed density per layer pair. Note that the proportion of independent pairs rapidly decreases, and thus increases for dependent pairs with graph depth.

### B. Control in Multitask Graphs

The characterization of different forms of dependencies above helps clarify the different roles that control can play in performance in multitask settings. First, we consider these qualitatively, and then use the framework introduced above to provide a formal analysis of the demands for control in such settings.

As discussed earlier, insofar as we are not concerned with reflexes, *some* allocation of control is required even for independent tasks, to “license” their execution by ensuring that activity in the corresponding output nodes is sufficient to overcome baseline inhibition (i.e., their thesholds for responding); that is, ∆*β* > *ω* + *β*. Control may be required at internal layers for similar reasons, and to augment processing where a pathway is weak (small *ω*). Thus, while tasks that are independent can be safely multitasked, they nevertheless require *some* allocation of control. Insofar as the analysis of control in such cases does not need to take into account any interaction among tasks, it can be viewed as an extension of the analysis in [13] to multiple independent tasks.

In contrast, for tasks that are either structurally or functionally dependent, control is needed not only to augment task processing, but also to mitigate the risk of conflict and ensure that the desired task(s) can compete with any stronger ones—that is, ones with larger *ω* values—with which they are dependent for which stimuli are present in the environment. This aligns with the role of control in neural network models, such as the model of the Stroop effect discussed above ([27, 28, 77, 86]. However, such neural network models, while mechanistically explicit, have been largely descriptive. Here, we use the information theoretic approach outlined above to provide a more rigorous formal analysis of the need for control, that provides the foundation for a normative analysis of the tradeoff between automaticity and control that we consider further on.

Using the expressions, and the multitask graph = 𝒰 (𝒱 ε^*l*^, 𝒫^*l*^,) introduced above, we can extend Eq. 18 to include the controlled efficacy of each task in a multitask ℳ as follows:

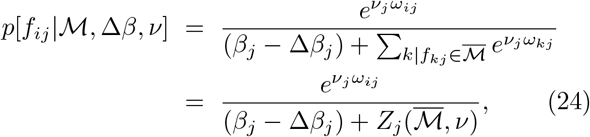

Here, ℳ specifies the intended set of tasks, whereas 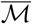 specifies the full set of tasks that may be engaged by the presence of functional dependence among the tasks in ℳ, and that can degrade their performance.

Note that Eq. 24 re-introduces the *v* control parameter, that appeared in the initial formulation of pathway efficacy (Eq. 15). We ignored it in the context of the single task graphs discussed above, as there it has effects that are qualitatively similar to (and trade off against) the effects of the **∆*β*** parameter. In the context of multitask graphs, however, there is a subtle but important distinction between the effects of **∆*β*** and *v*.

As discussed in Section II D, **∆*β*** regulates the overall responsivity of a node (by offsetting its inhibitory bias β), which is required in some amount to perform any task (except reflexes; see Section II A), and can be used to support the performance of ones that rely on weak pathways (or, as we discuss below, competition with tasks that rely on stronger pathways). In these respects, **∆*β*** can be thought of as reflecting a *selective* form of control, inasmuch as it licenses the execution of a specified task. This aligns with the assumption that the **∆*β*** parameters reflect the effect of learned control signals projecting to each processing node, mimicking the processing bias of task units in neural network models of control-dependent processing (e.g., [27, 79]).

In contrast *v* implements a strictly *modulatory* form of control, by modifying the sensitivity of the node to differences among its inputs. Critically, as *v* → 0, not only does the node become less discriminative, but it also becomes *more* likely to be active by chance (viz. with probability 0.5) irrespective of its **∆*β*** or ω parameters. [131] This would, of course, be deleterious when the **∆*β*** and/or ω parameters can be used effectively to implement a desired task. Under such conditions, increasing *v* can actually serve to *exploit* (i.e., further reinforce) their effects. However, for cases in which appropriate values of **∆*β*** and/or ω are *not* available to perform a given task (e.g., they have not yet been learned), *reducing v* can actually serve to increase the likelihood of performing that task (i.e., by chance). This can be thought of as supporting exploration, which can be valuable for discovery and learning [88–91]. Gain modulation of this sort has been proposed as a model of the effects of neuromodulatory neurotransmitters [92], and of norepinephrine in particular, in regulating the balance between exploitation and exploration [91, 93]. Since the timing of such neuromodulatory effects can be relatively precise [94, 95], although somewhat topographically diffuse [91], we assume that, while the *v* parameter can vary freely over time and can be modulated across nodes. It also important to note that the modulatory effects of ***v*** described above are relevant only in the context of multiple tasks because they describe a competition between tasks of different strenghts, at a given level of inhibition (**∆*β***), whereas when only one task is present the effect of ***v*** can be reabsorbed in the **∆*β***.

To consider how the control parameters **∆*β*** and *v* interact and can be optimized to perform multiple tasks, Figure 9 shows an example of a task graph for a network with a multitask combination ℳ composed of two pathways (shown in orange and green). In this case, the corresponding 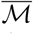 contains two additional mappings (shown as red edges) that introduce functional dependence and thus the potential for interference between the tasks constituting the desired multitask. In the case of strictly automatic processing considered above (see Eq. 18), the interference effects are determined fully by the ω_*ij*_’s for the pathways in 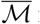 responsible for functional dependence among the tasks ℳ in to be performed. However, the ∆β and *v* terms introduced in Eq. 24, representing the effects of control, can be used to mitigate interference by modulating processing in the relevant pathways: These can be used to attenuate and/or eliminate processing along some pathways, restricting performance to only a subset of tasks or, in the limit, a single task within ℳ at a time. Based on Eq. 24, the control policy (i.e., set of **∆*β*** and *v* terms) that optimizes performance relative to strictly automatic processing can be determined, for a single-layered graph, as follows:

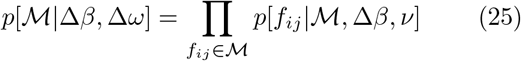

This can be further generalized to address a full multilayer graph, 𝒰, by calculating the product of the efficacies of each task pathway, *P* in ℳ and, once again, taking into account the extended multitask 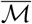 induced by any functional independence among those *P*’s:

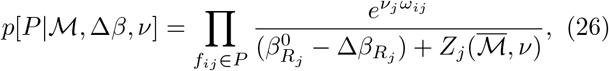

where *i, j* label tasks in *P* (mapping stimulus set *i* to response set *j*). The overall efficacy of ℳ is thus:

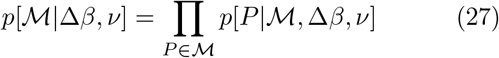

The corresponding multitask performance cost is then:

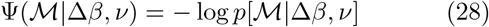

and the amount by which control and adaptive weights can modulate the interference in a multitask ℳ as:

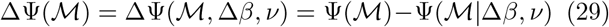

To illustrate how the expressions introduced above can be used to determine an optimal control policy for different graph configurations, we consider simple cases in which each task has only two possible values for the weights of the edges along its pathway, a strong one (ω=S) and a weaker one (ω=W). Specifically, we focus on multitasks involving functionally dependent tasks with two different patterns of pathway strengths: one in which the task pathways are composed of a mixture of strong and weak edges (SW; Figure 10a), and another in which all of the edges are similarly weak (WW; Figure 10b).

**Figure 9.**
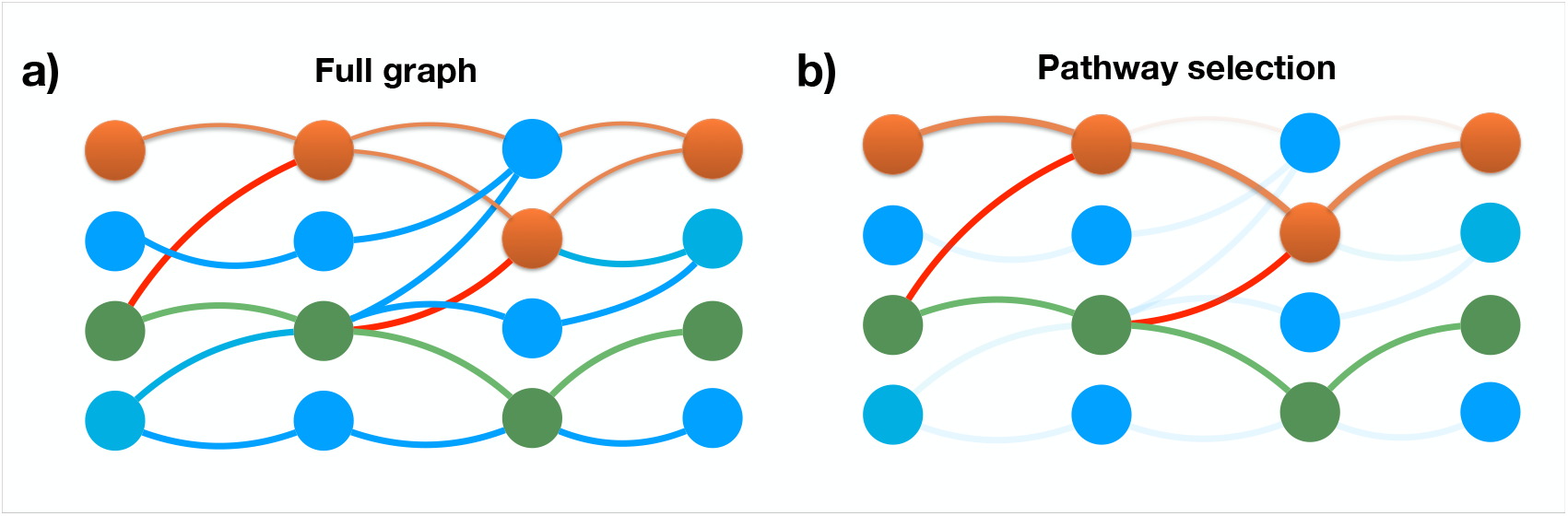
Pathway selection in a multitask graph. a) The full task graph for a multitask graph, 𝒰, showing the pathways (in orange and green) for two tasks in a specified multitask, ℳ, and the mappings for other tasks that introduce functional dependence between them (shown in red) and others that do not pose a risk of conflict (shown in blue). b) The pathways in the graph relevant for the multitask ℳ constituted by the two highlighted paths (orange and green, respectively) plus the edges in red, which are included in 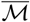 due to the presence of functional dependence and irrelevant pathways (faded).

**Figure 10.**
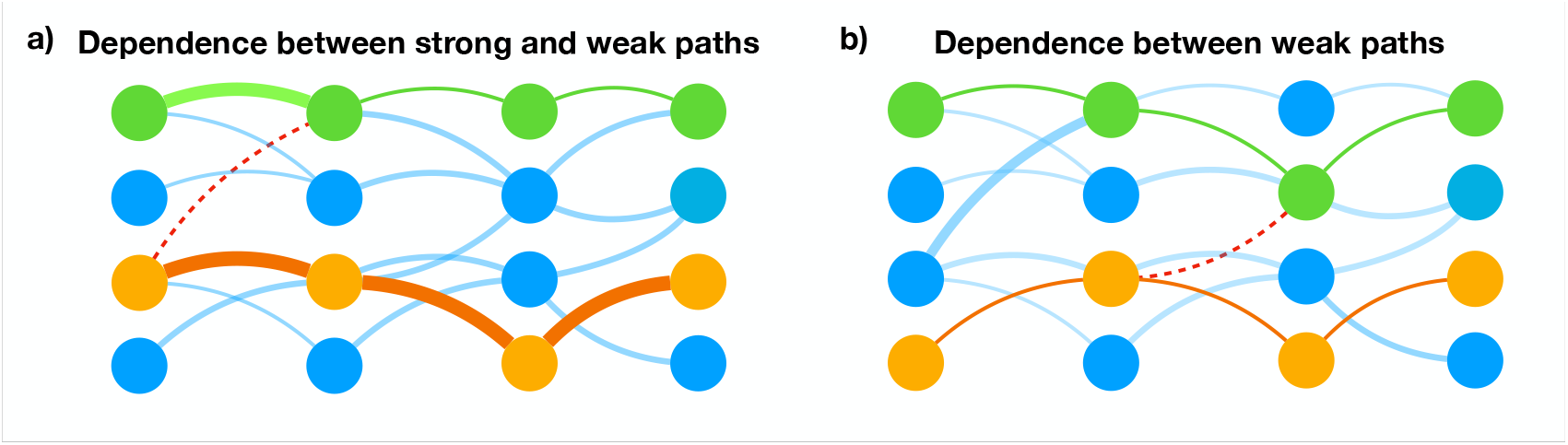
Examples of multitasks involving dependent tasks with different patterns of pathway strengths. Two multitask configurations, ℳ, each involving two tasks (orange and green pathways): a) the two pathways are composed of a mixture of stronger and weaker edges (SW in text), or b) the two pathways are composed of edges all of which have weak weights (WW in text). In both cases, the two tasks in ℳ are functionally dependent as result of the another pathway (red dashed edges) constituting 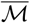.

Figure 11 shows the automatic processing costs Ψ (dark points) and controlled processing costs Ψ^∗^ (lighter points) for a large sample of pair-wise multitasks for the WW and SW cases in graphs of varying depth (from 3 to 8). Controlled processing costs were computed by optimizing the parameters (β, *v*) to minimize cost. As expected, allocation of control, improves the efficacy of processing by reducing performance costs: these are consistently lower in the controlled than the automatic condition in graphs of all sizes for both the WW (Figure 11a) and SW (Figure 11b) types of task pairs. Interestingly, the Ψ’s for multitasking pathways with similarly weak weights (WW, 11a) are much greater (performance is worse) than for ones with a mixture of strong and weak weights (SW, 11b). Nevertheless, even in the conditions showing the best performance (SW in graph with 3 layers), in no case can Ψ^∗^ be brought to zero. This is the result of functional dependence among the pathways; that is, the interference associated with functional dependence can be mitigated, but not entirely eliminated by control [132].

**Figure 11.**
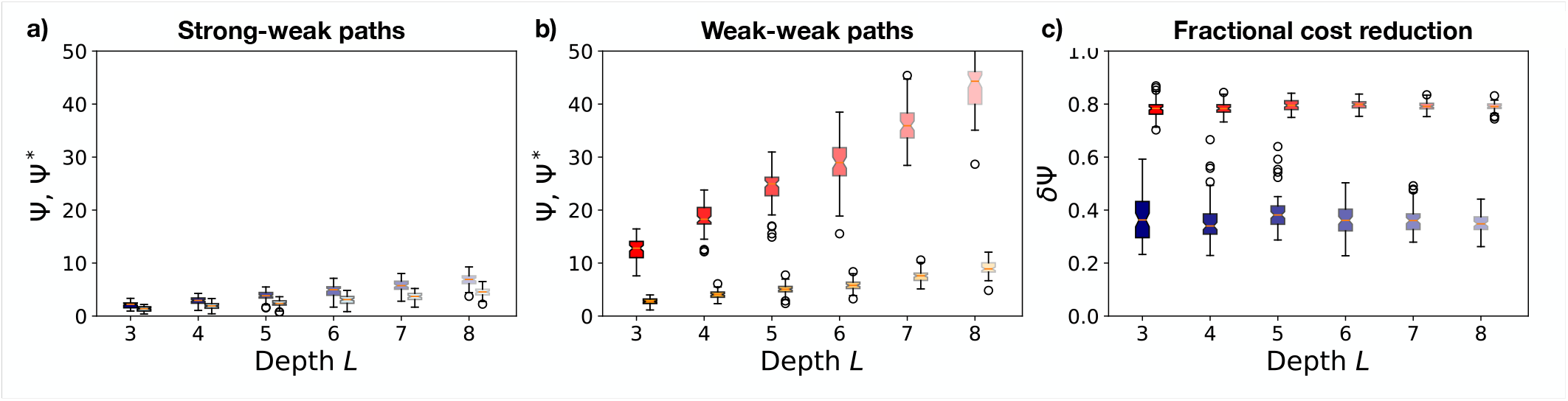
Performance costs for multitasks involving tasks with different relative strengths. Performance costs (automatic: Ψ; control-optimized: Ψ^∗^) for multitasking pairs of tasks chosen to have: a) both strong and weak pathways (SW; see example in Fig. 10a); and b) only weak pathways (WW; see example in Fig. 10b). In each case, performance costs are shown for strictly automatic execution (darker points) and control-optimized execution (lighter points). Note that, for graphs of all sizes, costs are less (performance efficacy is greater) for SW pairs relative to WW pairs, both for automatic and control-optimized execution. However, c) the *fractional* reduction in performance cost (i.e., improvement in efficacy) as a function of control, given by Eq. 30), is considerably greater for WW (violet and light blue) as compared to SW pairs (SW; red and orange), although SW pairs exhibit greater variance in this effect.

It is also interesting to note that, even though multitasking performance is overall better for SW pairs (including in the controlled conditions), the *fractional reduction in performance cost (i*.*e*., *improvement in efficacy* achieved by the allocation of control, calculated as:

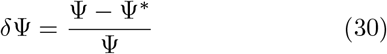

is substantially greater for WW pairs, as shown in Figure 11c (red points consistently higher than violet ones). This suggests that, although similarly weighted pathways tend to interfere more, their dependency can be modulated effectively by control, and that this holds irrespective of graph depth (Kruscal *s* = 35.1, *p* < 10^−3^ for WW, Kruscal *s* = 10.1, *p* < 10^−3^ for SW), although the variance in δΨ is greater for SW pairs (Levene test on paired distributions (F > 19 df 1 = 2, df 2 = 60, *p* < 10^−4^ for all comparisons) and seems to decrease with graph depth. These effects can also be seen in the distributions of control parameters for the various configurations, shown in Figure 12: for WW pairs, the optimized ∆β^∗^ values are distributed toward the upper limit of the range allowed, presumably because they have a substantial impact on cost; whereas for SW pairs they are smaller and more heterogeneous. Conversely, the *v*^∗^ values for the WW cases tend to take extreme values (0 and 2), while for SW we find a finer modulation of values between 1 and 2. This suggests that optimizing control for SW cases requires identifying more complex configurations of control signals (with greater information), in that SW cases require finer-grained adjustments to optimize performance, compared to WW cases, in which control is allocated more in a more binary (all-or-none) fashion, splitting between high *v*^∗^ values that amplify the effects of control, and low values that relegate the outcome of performance to pathway strengths in the absence of control (i.e., the effects of automaticity).

**Figure 12.**
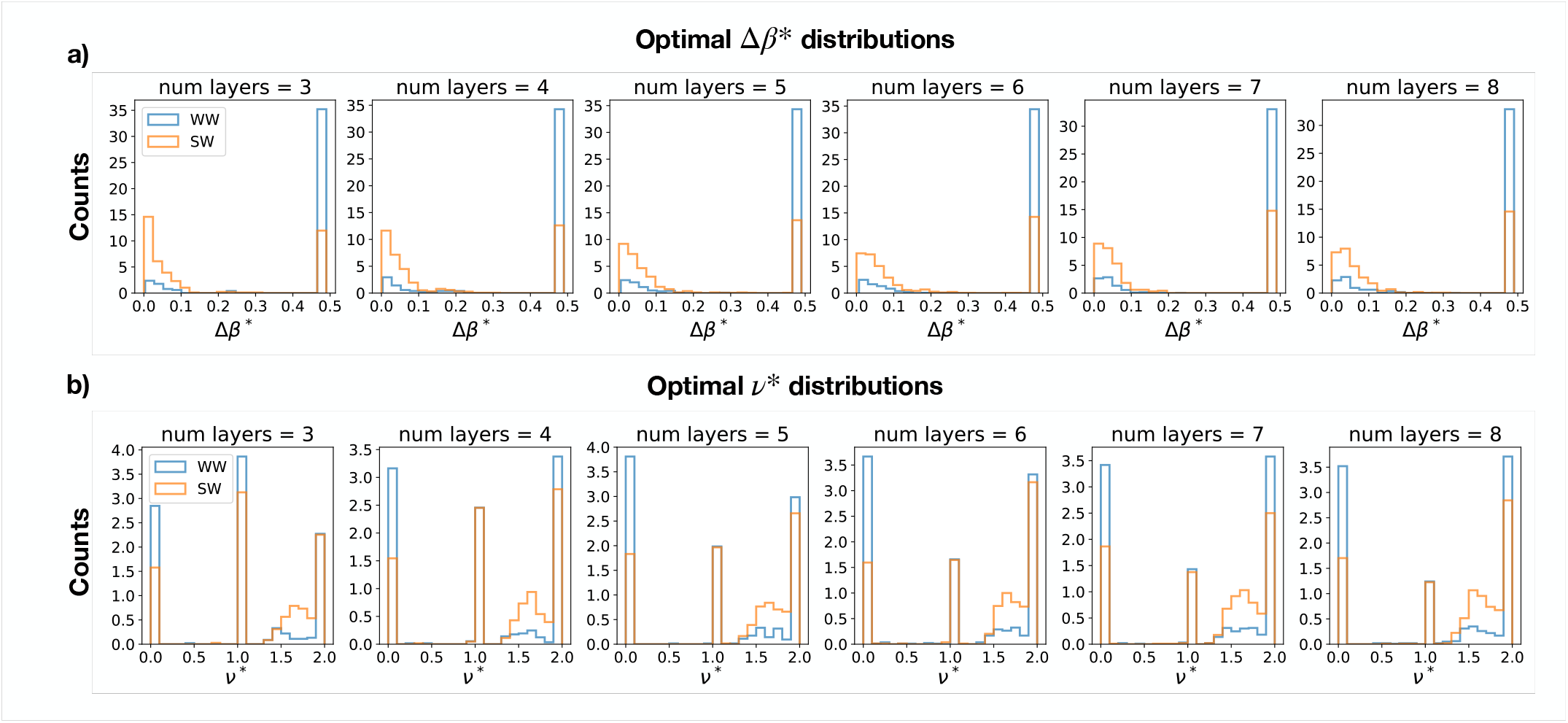
Differences in optimal control policies. Distributions of optimal control parameter sets *{*Δ*β*^∗^*}* (panel a) and *{ν*^∗^*}* (panel b) for the WW and SW cases.

## IV. THE TRADEOFF BETWEEN AUTOMATIZATION AND CONTROL

The framework introduced above permits quantification of the efficacies of automatic and controlled-based performance for sets of tasks given a neural network architecture used to perform them. The formulations we described all held the {***β*}** (intrinsic inhibition) and {***ω*}** (connection weight) parameters constant under the assumption that, to the extent that these are modifiable, they change at a much slower rate than the control parameters, {**∆*β*}** and {***v*}** . This is consonant with the widely held assumption that mechanisms of control are responsible for the short-term flexibility of human behavior. However, humans are of course also capable of remarkable flexibility over longer terms as well, including the ability to dramatically improve performance on tasks through practice, transitioning from what was initially a laborious and inefficient form of control-dependent processing to more fluid and automatic forms of processing. For example, a person who has never used a keyboard can quickly begin to type using one finger at time. However, while this exhibits remarkable flexibility (e.g., any finger can be used, and anything can be typed), it is also inefficient, something that can be overcome with sufficient time spent learning to touch type.

As discussed in the Introduction, the reliance on control for newly acquired skills, and the transition from control-dependent to automatic processing with practice, are some of the most widely recognized and studied phenomena in cognitive science [1, 40]. Various models have been proposed that provide a mechanistic account of this transition [21, 27, 97, 98], including recent work that has begun to address this from a strategic perspective [49, 50]; that is, how and when people decide to invest the time and effort to transition from control-dependent to automatic processing? The framework outlined above lays the foundation for a mathematically rigorous, normative answer to this question. In addition to addressing how control may be optimized for performing a set of tasks that are dependent on control (as formulated in the preceding section), it also allows us to determine whether and when the system should exploit the immediate benefits of control-dependent processing (through rapid adjustment of {**∆*β*}** and {***v*}**), at the cost of efficiency due to functional dependence, or seek to improve efficiency by restructuring the graph to strengthen and/or develop task-dedicated pathways that afford parallel processing (through the slower adjustment of {***ω*}**), but at the cost of the time and effort required by the training to do so.

Below, we formalize this optimization with respect to a given graph architecture, a set of tasks to be performed, a finite horizon over which reward is to be optimized, and the relative speed with which the {**∆*β***} and {***v***} versus {***ω***} parameters can be adjusted.

### A. Tradeoff Between Current Serial and Future Parallel Execution

The optimization problem outlined above can be framed as one of finding the control policies and/or investments in automatization for a given set of tasks, over a specified period of time, that maximize total cumulative reward (or, alternatively, average reward rate). For a given graph, the set of tasks to be performed can be considered as a multitask, ℳ, that must be performed in the context of an extended multitask 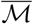 that includes any other tasks defined by the graph’s architecture that induce functional dependence among the tasks in ℳ. Of course, at any point in time, the environment and/or internal needs of the agent may render some subsets of tasks in ℳ more viable and/or desirable than others (e.g., depending on whether the relevant stimuli are present, actions are possible, and/or results are presently valued); those subsets, in turn, will be associated with different extended multitasks. These are critical factors for addressing the complexity of any real-world scenarios and agents, an issue to which we return in the Discussion.

Here, as a tractable starting point, we assume a stationary environment in which all tasks are uniformly viable and desirable. Accordingly, executing as many as possible, as reliably, and as frequently as possible will yield the greatest rewards. However, as the preceding analyses make clear, the extent to which this can be achieved depends on the graph architecture—that is, its {***β*}** and {***ω*}** automaticity parameters. In the extreme, if all tasks are independent, they can all be performed in parallel, and the only constraints on reward rate are the environment and the speed with which the tasks can be performed—that is, the absolute values of the automaticity parameters for each task [133]. Conversely, if all pairwise combinations of tasks are either structurally or functionally dependent, then only *one* can be performed reliably at a time; that is, performance is constrained to be serial, and the reward rate is directly proportional to the number of tasks in ℳ. What is less clear is the optimal policy for graph architectures that fall somewhere between these two extremes; that is, in which some combinations of tasks can be safely parallelized, but others cannot. For such a graph structure, and a set of tasks to be performed, the question is: How can those tasks best be combined into multitask subsets, such that the tasks within each subset are as independent as possible (i.e., have high multitasking efficacy) and can thus be executed effectively in parallel, while the subsets are executed in series to avoid conflict among tasks that are dependent on one another. Below, we formalize this partitioning, and the overall efficacy of performance it yields, and use this to quantify the best performance that can be achieved for a specified set of tasks in a given graph architecture. In the section that follows, we then show how this can be used to address the tradeoff between automaticity and control, by comparing the current performance efficacy of a graph against the benefits of improving its efficacy by modifying its structure to diminish dependence among the specified tasks, weighed against the costs (in time, and thus average reward rate) of doing so.

#### Optimization of multitasking covers

The optimal execution of multiple tasks may be achieved by varying the degree of serialization in a continuous manner, from time step to time step [54]. To ease formal analysis, we consider cases in which an agent either fully serializes or parallelizes tasks for execution. We begin by defining a multitask ℳ containing tasks 𝒯 to performed, and seek a cover, Θ, of the tasks 𝒯 in ℳ, that divides them into subsets Θ_*i*_ such that Θ_*i*_ ∩ Θ_*j*_ = ∅ for all i, j and the union ∪_*i*_ Θ_*i*_ = *T* . Denoting Θ_*i*_ = |Θ_*i*_|, we have that Σ_*i*_ Θ_*i*_ = Θ. Thus, the cover Θ corresponds to a collection of subsets of tasks (Θ_*i*_’s) formed by dividing all of the tasks in ℳ into *s* disjoint subsets (with the index s starting at 1, as there is always at least one set in Θ), such that every task in ℳ belongs to one and only one Θ_*i*_. The tasks in each Θ_*i*_ are considered a multitask (i.e., they are executed at the same time), while the Θ_*i*_’s will be executed serially. Accordingly, the goal is to construct Θ such that overall processing efficacy is maximized, by maximizing the multitasking efficacy for each subset Θ_*i*_ (given by the automaticity parameters of tasks in the subset, and the optimal control policy for that multitask, as described in Section III) and minimizing the total number of Θ_*i*_’s to reduce serialization costs and thereby maximize reward rate.

The overall efficacy of a cover Θ can be expressed as:

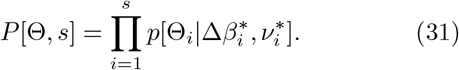

where the ^∗^ represents optimized parameters. Furthermore, we can express the reward rate associated with Θ as:

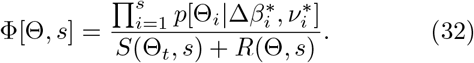

in which the denominator describes the overall time it takes to execute the full set of Θ_*i*_’s in Θ. The first term in the denominator, S, includes the number s of sets in Θ that contribute directly to the serialization cost. S also includes two additional terms: 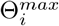, which denotes the execution time of the slowest task in each of the Θ_*i*_’s; and R(Θ, t), that denotes the total time required for switching between performing different task subsets (Θ_*i*_’s), which may involve both the regulation of control parameters [6, 46] as well as passive processes associated with the updating of task sets [5, 31]. Thus, these terms reflect the aggregated effects of dynamics of performance of individual tasks (Θ_*t*_ in *S*) and switching between subsets of tasks (*R*(Θ, *s*), respectively. Although our preceding analysis of performance for individual tasks and multitasks ignored these effects (see Note V D), here we include them as we are now interested in the reward *rate* associated with the performance of *sequences* of tasks and/or multitasks. Thus, while the effects for individual tasks or multitasks may be relatively small, they may accumulate over sequential executions.

Note that the efficacy *P*[Θ, *s*] can be re-written explicitly in terms of accuracy, by replacing *p*[Θ_*i*_ | ∆*β*_*i*_, *v*_*i*_] in Eq. 31 with 1 − ER(Θ_*i*_), where ER is error rate. Accordingly, Eq. 32 can be expressed in the more familiar form of a reward rate [37, 100]:

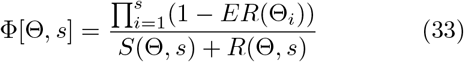

Thus, maximizing Φ(Θ, s) is equivalent to maximizing reward rate. Φ(Θ, s) in Eq. 33 can also be re-expressed in terms of costs, similar to Ψ, as:

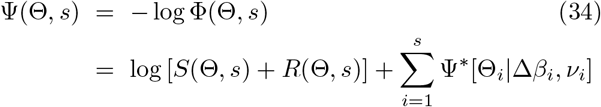

This is useful for considering Φ in graph theoretic terms, that in turn, provides a useful approach to the optimization problem. For example, an ideal cover, Θ, would be composed exclusively of Θ_*i*_ each of which is a subset of tasks that are independent of one another, constituting multitasks that can be performed entirely in parallel without any allocation of control. In that case, Eq. 34 can be reduced so that Φ[Θ, s] depends only on the serialization and switch costs:

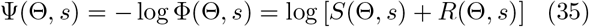

A cover comprised exclusively of subsets of tasks that are independent of one another in the interference graph for the task graph represents a *coloring* [101], and thus the optimal Θ corresponds to a minimal graph coloring of the interference graph for ℳ. Previous work [19] reports an approach to solving this problem. This formal treatment of partial task serialization relates closely to the optimal scheduling of task processes in symbolic processing architectures [83, 84].

*Example*

As an illustration of how the formalisms introduced above can be used to find the optimal cover for a given graph configuration, Figure 13 shows the results of numerically optimizing Φ over multitasks comprised of three tasks with different types of pathways and in graphs of different depths, taking into account both the costs of control and serialization. Graphs and task sets of each type were constructed and, for each, Eq. 33 was used to compute Φ for each of the three possible types of cover Θ for the three tasks in multitask ℳ: a single set, such that all three tasks are performed simultaneously (3 *tasks*); two Θ_*i*_ subsets, in which two of the tasks are performed simultaneously and one on its own (2 + 1 *tasks*); and three Θ_*i*_ subsets, each containing a single task, so that the tasks in ℳ are all performed serially (1 + 1 + 1 *tasks*). The plots in Figure 13 show the frequency of each type of cover for different graph depths and types of task combinations. Note that as graphs become deeper, and thus the likelihood of dependencies increases (Fig. 8), the optimal serialization strategies shift from full multitasking (3 *tasks*) to intermediate (2+1 *tasks*) or full serialization (1 + 1 + 1 *tasks*). Similarly, sets composed of weaker tasks (13b) are associated with a shift toward greater serialization as compared to those with strong tasks (13b), reflecting both increased processing costs and the greater likelihood of interference with multitasking.

**Figure 13.**
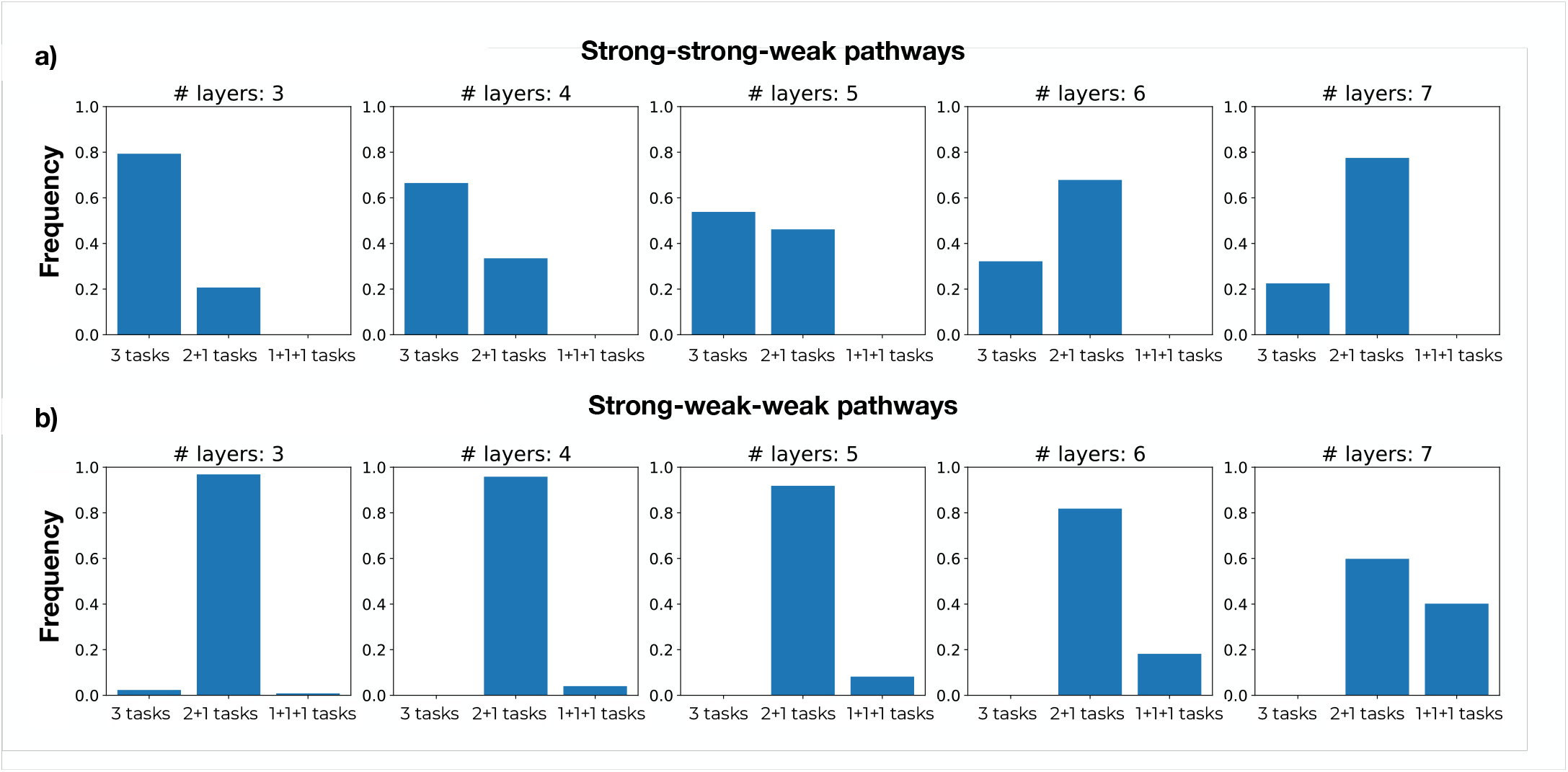
Optimal serialization for three tasks. Frequency of types of partitioning of three tasks into subsets (Θ_*i*_ covers) to minimize overall performance cost Ψ (i.e., optimize efficacy) in graphs of various depths (3-7 layers), in which the balance of strength among the three tasks is varied in a manner similar to Figure 12: a) two strong and one weak; or b) one strong and two weak. Each plot shows the frequency with which each type of partitioning of the three tasks (cover Θ_*i*_) yields the best Ψ: all tasks in a single set and thus executed in parallel (3 *tasks*); two tasks in one subset executed in parallel, and in series with the remaining task executed on its own (2 + 1 *tasks*); or all three tasks executed in series (1 + 1 + 1 *tasks*). Results were obtained from 1000 graph instances for each depth, with randomly assigned weights and sets of tasks of each type chosen from them.

### B. Joint Optimization of Control and Automatization

The optimization described in the previous section sought to partition a set of tasks in a multitask ℳ into a cover Θ comprised of one or more subsets of tasks that are as mutually independent as possible. The extent to which this is possible is constrained by the graph structure, including the relative strengths of the pathways for the tasks in ℳ, as illustrated in Figure 13. Alternatively, Φ (and thus reward rate) could be maximized for a given by ℳ modifying the graph structure itself; that is, by modifying the ***ω*** for the tasks in so as to make them more independent of one another, and thus more effectively executed in parallel. As discussed earlier, we assume that the ***ω*** can be modified over the course of learning, which occurs at a much slower time scale than adaptation of the control parameters, **Δ*β*** and ***ν*** parameters used to implement the Θ for a given ℳ. This formalizes the tradeoff between reliance on control-dependent processing and the acquisition of automaticity as an intertemporal choice: For a given time horizon and multitask ℳ, is it preferable to pay the cost of control-dependent processing (i.e., constrained Φ) for the current graph structure, or to pay the cost of increasing Φ by modifying the relevant *ω*s to afford better multitasking and thereby permit a better Θ (and therefore Φ). Which is preferable will, of course, depend on the probability distribution of the tasks in ℳ, and their corresponding reward values.

To formalize this tradeoff, we can refine the definition of Φ to include the structure of the graph at a given time *t* as *ω*(*t*), and Φ = Φ[Θ, *s, ω*(*t*)], which can change with time as a function of learning. Then, over a horizon of total time *T*, the cumulative reward is given as the integral:

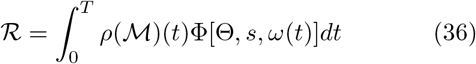

where *ρ*(ℳ)(*t*) is the probability at time *t* of encountering the task set ℳ, which the agent can perform optimally using the task cover Θ, and the average reward rate is 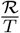.

Importantly, the total time *T* can be split into two contributions, *T* = *T*_*exec*_ + *T*_*learning*_. The first component, *T*_*exec*_, accounts for all the task execution times, and is, therefore, a function of: i) the distribution of tasks in the environment (*ρ*); ii) the pattern of (in)dependence among those tasks as determined by the temporally evolving ***ω*(*t*)** that determines both the optimal cover Θ; and iii) the corresponding serialization and reconfiguration times (*S* and *R*). The second component, *T*_*learning*_, accounts for the time required to adapt ***ω***, that results in a temporal trajectory of *ω* configurations 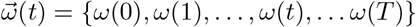.

Given this formulation, the strategy that maximizes *R* depends on two critical factors: the rate at which {***ω}*** s can adapt, and the time horizon *T* over which optimization is sought. The former is determined by the learning rate, that we assume is intrinsic to the system and/or task, while the latter is determined by the environment. In general, it is easy to see that both faster learning rates and longer time horizons will favor the acquisition of automaticity, as it provides more time to benefit from the investment in learning {***ω}*** s that maximize independence among tasks and thus permit parallel execution of task combinations in ℳ. However, whether this is the optimal policy (i.e., maximizes *R*) will depend on the actual learning rate[134] and time horizon. Furthermore, it will also depend on the value of continuing to rely on control-dependent processing, which provides more immediate reward, albeit less than what could be achieved in the same circumstances once automaticity has been achieved. For example, if the time horizon is sufficiently short relative to the learning rate, such that *T*_*learning*_ *< T*_*exec*_, then the investment in acquiring automaticity will not be worth more than the value of continued reliance on control (e.g., it may not be worth learning to touch type just to write a single essay). However, for longer time horizons, the relative values may reverse (e.g., if the goal is to make a living as a stenographer). That is, choosing between control-dependent processing versus the acquisition of automaticity is an intertemporal choice that is determined by the relative values of *T*_*exec*_ and *T*_*learning*_.

Previous work has described how the tradeoff between control and automaticity may be optimized in the context of specific learning algorithms and neural network architectures [49, 50]. The formulation outlined above provides a more general framing of the problem. While a full solution may be demanding, and potentially intractable for realistically complex conditions, below we illustrate how, with some simplifying assumptions the formulation can be used to analyze the trade-offs between control and automaticity in order to maximize *R*. This may serve as an example of how natural agents, using heuristics, may make similar computations, and how artificial systems might be designed to do the same.

#### Automatization in the extended Stroop task

As an illustration, we consider the case of the extended Stroop task (see Figure 2b), in which word pointing and color naming are initially functionally dependent but, with practice, automaticity can be acquired for word pointing, allowing it to be performed in parallel (i.e., multi-tasked) with color naming. This can be accomplished by learning a new set of associations for word pointing that can be used to map word stimuli (*S*_2_) to manual responses (*R*_2_), but that are mediated by internal representations (shown as *H*_3_ in Figure 14a) that are distinct from those used for word reading (*H*_2_). That is, with learning, *H*_3_ can be used to strengthen a latent (weak) pathway [135] and/or form a new one (*T*_3_(*S*_2_ *H*_3_ *R*_2_) that is fully independent of the color naming pathway (*T*_1_ = {*S*_1_ *→ H*_1_ *→ R*_1_}, and thus allow word pointing to be performed in parallel. We assume that *T*_3_ starts with small initial weights (i.e., small ***ω*** s for *S*_2_ *→ H*_3_ and *H*_3_ *→ R*_2_), but that these can be made stronger with practice on the word pointing task as they are updated with learning rate λ.

**Figure 14.**
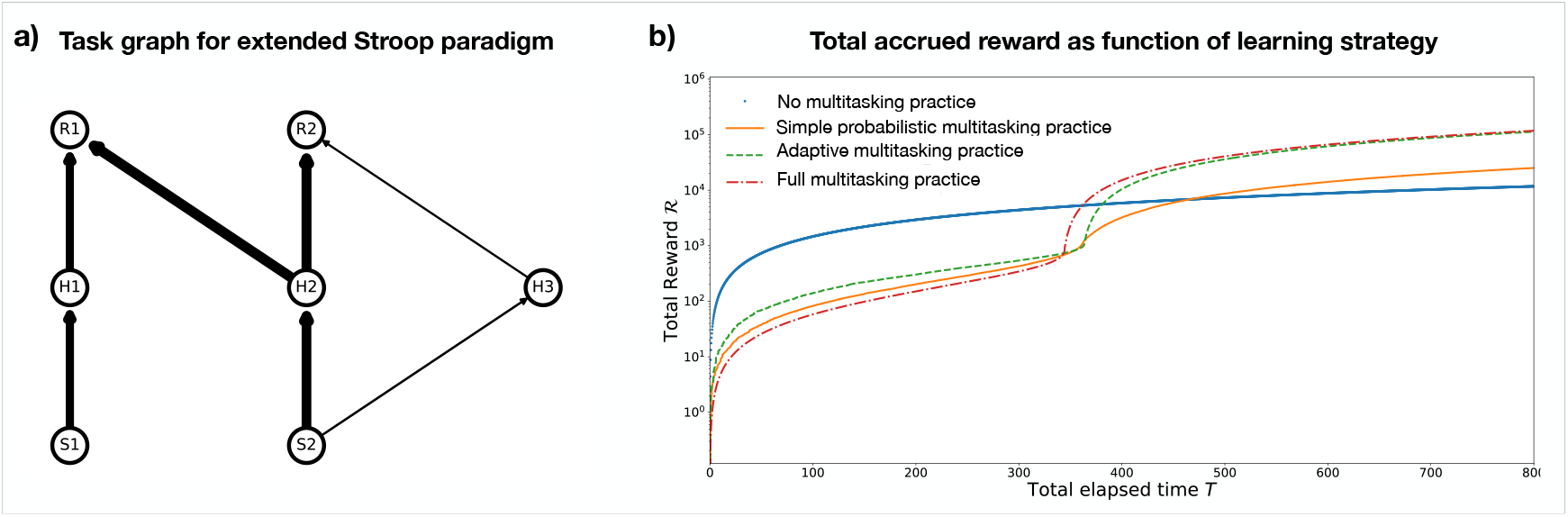
Total cumulative reward accrued under different automatization policies in the extended Stroop task and fixed learning rate of *λ* = .001. a) Task graph for the extended Stroop task (based on the graph shown in Figure 2b) in which color naming relies on *S*_1_ *→ H*_1_ *→ R*_1_ (*T*_1_) and word pointing on *S*_2_ *→ H*_2_ *→ R*_2_ (*T*_2_, with functional dependence induced by word reading due to edge *H*_2_ *→ R*_1_); but here with the addition of internal representation node (*H*_3_) that provides an alternative, initially weak pathway for word pointing, *S*_2_ *→ H*_3_ *→ R*_2_ (*T*_3_), that is independent of the color naming pathway *T*_1_. b) Plots of the total accumulated reward (*R*) over executions (proportional to *T*) for different practice policies (see text for description). Note that *R* is expressed in log units, and thus linear accumulation of reward (e.g., in the *no practice* condition) follows a standard logarithmic form.

To quantify this, we consider the case in which the agent is confronted with repeatedly performing the color naming and word pointing tasks, and thus performing either {*T*_1_, *T*_2_}(constrained by serial performance) or {*T*_1_, *T*_3_}(which permits parallel performance, but must be acquired through practice). Figure 14b shows the total accrued reward ℛas a function of time (i.e., number of executions of the two tasks) under four policies that vary with respect to whether and how {*T*_1_, *T*_3_}is “practiced:”

- *no multitasking practice*: this serves as a baseline, in which learning rate λ = 0, performance of the word pointing task relies exclusively on the existing set of word representations (i.e., pathway *T*_2_), and word pointing and color naming continue to rely on control for serial execution of {*T*_1_, *T*_2_}(blue solid line).
- *simple probabilistic multitasking practice*: {*T*_1_, *T*_3_}is performed with a probability *p* (i.e., executing word pointing using *T*_3_ in parallel with color naming is practiced with this probability, in this case we adopted *p* = 0.2). Although performance is poor early on, *T*_3_ is strengthened with each execution, so that performance gradually improves, and eventually, cumulative reward exceeds that of strict reliance on control (orange solid line).
- *adaptive multitasking practice*: {*T*_1_, *T*_3_}is also performed probabilistically, but adaptively as an inverse function of its cost (Ψ) relative to that of {*T*_1_, *T*_3_}:

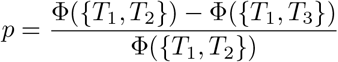

This policy is superior to the simple probabilistic practice, and converges to the same total reward as full practice (see below), but with a different tradeoff, yielding a higher initial reward but deferring the benefits of automaticity (green dashed line).
- *full multitasking practice*: {*T*_1_, *T*_3_}is executed on every trial. This yields the lowest immediate rewards but the earliest benefits of automaticity (red broken line).

In general, Figure 14 shows the qualitative pattern of intertemporal choice: over short temporal horizons it is better to forgo automatization and exploit the more immediate benefits of control-dependent processing by relying on {*T*_1_, *T*_2_}, even though this involves slower serial execution; whereas for sufficiently long temporal horizons, the deferred benefits of investing in automatization pay off, yielding overall greater cumulative reward. Importantly, the figure also shows that formalizing the processes involved can reveal subtler, quantitative effects associated with different policies that may be relevant to strategic decision-making as a function of differences in horizons and temporal discounting, that we consider in the next section.

#### Effects of online estimation and discounting

Thus far, we have considered the total cumulative rewards obtained under different automatization policies from the point of view of an “ideal observer” (i.e., with full knowledge of all relevant factors such as learning rate and task likelihood), using a fixed policy and, critically, assuming an infinite time horizon with no temporal discounting of future reward. However, in realistic scenarios, it is likely that the agent can extract estimates of its efficacy and reward rate based on its preceding experience with the task, and how these affect improvement, which it can use to generate expectations for the effects of practice on further improvement (e.g., priors on performance based on effects experienced in the past), and to take account of the temporal horizon *T*_*h*_ it has available for such practice, subject to discounting the value of future rewards. To capture these effects, we assumed that the agent could estimate future reward based on the learning curve (a logistic function relating the past amount of practice to the rate of improvement in performance).

To consider an agent that chooses a policy at a given point in time *t* based on these factors, we can formulate a simple estimate of what, at that moment, it can expect to accrue by sticking to a policy *P* until a horizon time *T*_*h*_ as:

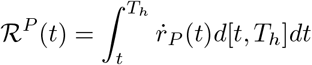

where *r*_*P*_ (*t*) is the estimated reward rate for policy *P*, and *d* is an exponential discount function *d*(*x*) = *e*^*−αx*^ for some discount factor *α*. Given multiple policies 𝒫= {*P*_0_, *P*_1_, … *P*_*k*_ }, the probability that the agent chooses a certain policy *P*_*i*_ at time *t* can be defined as:

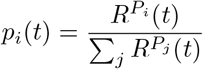

Figure 15 shows the relative choice frequency between two of the policies outlined above — i) serial execution of *T*_1_ and *T*_2_ (no multitasking practice, as a baseline), and parallel execution of *T*_1_ and *T*_3_, with updating of the weights of *T*_3_ over the course of learning, as described ii) above (adaptive multitasking practice) — for four horizon times (*T*_*h*_ = 0, 20, 100, 500, where 0 corresponds to the situation without any learning) and three discount rates (*α* = 0, 0.2, .5, where 0 corresponds to no discounting of future value). In each case, the agent repeatedly executed the two tasks, each time selecting between the two policies based on the average of instantaneous reward rates obtained in previous execution of the same policy and estimated future value of the policy based the effects of practice experienced to that point, for the specified *T*_*h*_ and *α*. The effects are consistent with those observed in Figure 14b, with the value of automatization increasing for both longer horizons (larger *T*_*h*_) and lower discounting of future value (smaller *α*) for a given learning rate.

**Figure 15.**
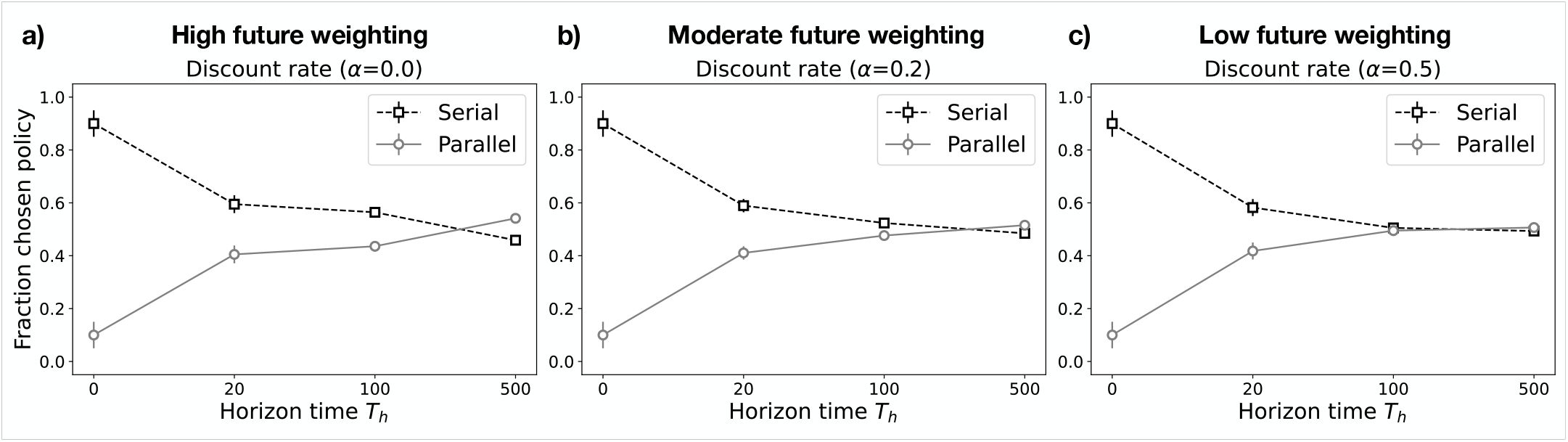
Choice of execution policy for different time horizons and discount rates, and fixed learning rate of *λ* = .001 (same as used in Figure 14). Frequency of execution of serial policy (*T*_1_ and *T*_2_ in series) versus frequency of execution of new parallel pathway (*T*_1_ and *T*_3_) (see text for description) for an agent for three possible horizon times (*T*_*h*_ = 0, 20, 100, 500) and three discount rates (*α* = 0, 0.2, 0.5, respectively panels a-b-c). For longer horizons the preferred policies shift toward automatization (i.e., parallel execution of *T*_1_, *T*_3_), but this interacts with discount rate, such that automatization is favored to a greater extent when future value is discounted less.

The observations suggest that an agent that favors future value (low discount rate) can, under appropriate circumstances (availability of long time horizons for practice) favor the development of separated, task-dedicated representations in order to parallelize performance (i.e. develop automaticity). At the same time, they also suggest that a large range of parameters favor the use of reliance on shared representation (at the expense of less efficient serial processing), which may be consistent with a general bias toward control-dependent processing when confronting the need to acquire a novel task. While this is of course contingent on learning rate, there may also be a strong bias toward lower learning rates to avert the well-known problem of catastrophic interference in neural networks associated with higher learning rates [34, 104]. More generally, it illustrates the usefulness of the formulation, and invites its application both in analyzing empirical findings concerning human performance, and building artificial agents that can autonomously optimize the tradeoff between the flexibility of “general purpose” computing and the efficiency of tailoring processing to specific tasks. We return to a broader consideration of this issue in the Discussion below.

## V. DISCUSSION

### A. Summary

In this article, we introduce an information-theoretic approach to quantifying the cost of control in neural network architectures. Our approach builds on an understanding of what has come to be considered, from a cognitive and neuroscientific perspective, a fundamental role of control: managing the risk of conflict associated with shared representations [21, 27, 28]. Critically, this can be overcome through additional training, by learning to separate the representations responsible for conflicting tasks, thus permitting parallel processing (i.e., multitasking) [9, 21]. These observations point to a fundamental tradeoff between:

1. the efficacy of learning and flexibility of processing (via generalization) afforded by shared representations, but at the cost of the dependence on control to serialize processing;
2. the efficiency of automatic, parallel processing (e.g., multitasking) afforded by separated, task-dedicated representations, but at the cost of the additional time and effort required for training to develop these [12].

This formulation of the relationship between control-dependent and automatic processing has been used to account for a wide range of behavioral phenomena associated with human performance [21], and motivated formal analyses of the capability of neural network architectures that may be relevant both to understanding human performance as well as the design of artificial systems [19, 32, 50].

Here, we extended, and lent additional rigor to these ideas, by casting them in formal terms that allowed them to be integrated into an information-theoretic framework developed in prior work to analyze the effects of cognitive control [13]. That work focused on the performance of *individual* tasks, and the role of cognitive control in augmenting processing to ensure their reliable execution. Here, we extended the framework, by relating it to more fine-grained analyses of the neural network architecture required to perform an individual task (e.g., the patterns of performance, including the dynamics [see A] in response to individual stimuli and responses), as well as to include *interactions* among tasks in multitask graphs, and used it to address: the role of control in augmenting weaker tasks when in competition with stronger ones; how performance can be optimized (operationalized as maximizing reward rate) by selecting which tasks to execute at any one time; and how total cumulative reward can be optimized through decisions about when to rely on control for serial execution versus investing the time and effort in automatization to achieve the efficiency of parallel processing.

Central to the approach is the notion of *processing cost* or, inversely, *processing efficacy*. We first demonstrated that dependencies between tasks (in the form of shared representations) impose constraints on the processing efficacy of a neural network by limiting the number of tasks that can be performed at once, and showed that such limitations scale with the number of processing layers in the network, and are remarkably resistant to increases in the size of the network. We then provided a formal analysis of optimal efficacy for a given task graph structure and set of tasks to be performed. Finally, taking account of the fact that optimal efficacy can be improved by reconfiguring the graph over a longer time frame through learning, we formalized the tradeoff between the allocation of control in an existing graph and the investment in learning to reconfigure it, in terms of the joint of maximization of cumulative reward over a specified horizon, through the optimization in the allocation of control and the acquisition of automaticity through learning. This framework provides a foundation for analyzing how neural network architectures—whether biological or artificial—may manage the tradeoff between control-dependent and automatic processing, given the relative benefits and costs of each and how these relate to the task environment.

### B. Relationship to Previous Work

#### Information theoretic work

As noted above, the work presented here is grounded in prior work introducing an information-theoretic framework for analyzing the effects of cognitive control [13]. According to this approach, cognitive control serves to integrate information about a stimulus and task context into a task-dependent response [13]. While this approach offers a formal distinction between controlled and automatic processing for single tasks in information-theoretic terms (with the former but not the latter integrating information about the task context), it does not consider dependencies between multiple tasks. Zénon et al. [18] have proposed a more fine-grained analysis, according to which the information integrated in the service of one task—forming a posterior distribution of responses given the stimulus and task context—can influence the prior distribution of responses for the same stimulus in the context of another task, and thus the amount of information to be integrated for the other task. However, their approach does not take into account the representational structure of those tasks; that is, the extent to which the tasks interact with one another as a consequence of shared representations. By considering the implementation of tasks in a neural network architecture, our treatment takes account of dependencies between tasks induced by shared representations. This allows us to formulate, in information-theoretic terms, the effect of these dependencies at different levels of processing (from the processing of individual stimuli and responses within a single pathway at the neural network level to the interactions among tasks at the graph level), as well as the influence of factors intrinsic to the processing mechanisms in neural architectures, such as inhibition, the modulatory effects of control, and the depth of the network.

#### Graph theoretic work

The work presented in this article also extends previous graph theoretic analyses of the multitasking capabilities of neural networks, that were restricted to unweighted graphs in which interference and control are treated as all-or-nothing effects [19, 32, 105]. Here, we exploited the information-theoretic formulation to consider these as graded effects, and cast the tradeoffs among parameters as optimization problems. However, the formulation presented here continued to treat the task-specific mapping of individual stimuli to responses as having uniform strengths. This may be a reasonable assumption on average (e.g., the strength of the mapping from the word RED to the utterance “red” is roughly comparable to that from GREEN to “green”), but is certainly violated in some contexts (e.g., in a task that mapped red to stop, green to go and blue to wait). This might be usefully addressed by integrating the framework presented here with recent work on the relationship of control to statistical learning and semantics [79].

#### The tradeoff between the allocation of control and the acquisition of automaticity

As noted in Section III, previous work has begun to address this question both analytically [49] and in deep learning architectures [50]. However, these have involved highly abstracted forms of the problem (e.g., using simplified assumptions about learning curves, categorical costs, and Bayesian optimal estimation) or implementations in deep learning networks using more complex, realistic environments (e.g., video images as inputs) but restricted numbers of tasks. For example, Sagiv et al. [49] contrasted two pre-specified, extreme task configurations involving either quickly learned shared representations and strictly serial processing (with a fixed, pre-determined serialization cost), or more slowly acquired separated representations and fully parallel processing. The analysis of this system demonstrated that over a wide range of parameters (e.g., relative learning rates and time horizons), control-dependent, serial processing was favored over the acquisition of automaticity, but that this can reversed under reasonable conditions (e.g., more severe serialization rates or longer time horizons). Similar results have been obtained using a deep learning network trained in a more complex but realistic environment (with video images as inputs, and object recognition, object localization, and navigation tasks) [50]. The work presented here helps bridge these lines of work, by providing a framing of the problem that can take account of the complexities associated with neural network implementations, including standard learning algorithms and dependence among tasks as continuous and heterogeneous factors, while in a form that is more tractable to formal analysis than the full implementation of a neural network model.

#### Task scheduling in symbolic models and serial versus parallel processing

Dependencies between individual tasks and concomitant requirements for serial processing have been considered extensively in the context of scheduling the execution of tasks in symbolic (e.g. production system) architectures, such as EPIC [106] and threaded cognition [84]. However, to our knowledge, these have not provided a formally rigorous, normative analysis of the problem. Furthermore, they do not take into account the graded effects of representational sharing that are a critical source of constraints on multi-tasking and the requirements for control in neural network architectures, and that are accommodated in the information-theoretic framework we have described here. Rather, in symbolic models, constraints on multitasking are generally assumed to arise from the limited capacity of a central executive responsible for scheduling and the allocation of control, and/or a set of scheduling rules inherent to the processing architecture [84] that decide which resources get allocated to which tasks. It is worth noting, however, that this approach *has* been used to address empirical observations, and interpret these as evidence that humans shift between different processing modes (serial versus parallel task execution) depending on characteristics of the task environment [107]. At the same time, others have suggested that serial versus parallel processing may be better viewed as lying along a continuum, depending on the degree of interference between processing pathways [54, 108], a view that is more closely aligned with neural network models [21], and the framework presented here. While we have not explicitly considered intermediate solutions, they may be derived from this framework by considering micro-dynamics of processing within the processing nodes. While we have simplified our formulation by abstracting over such dynamics, in Appendix A we should how our framework can be related to previous work that addresses how these dynamics are critical for reward rate optimization in neural systems [47, 48, 109–112]. Such micro-dynamics also remain crucial for addressing the temporal dynamics of interference within a task [77, 78, 113], as well as between-trial dynamics of control allocation to a single tasks [28] and across multiple tasks [29, 46–48].

### C. Simplifications and Future Work

The formalisms introduced in this article rely on several important simplifications that should be addressed in future work.

#### Interference versus facilitation

First, analyses assumed that information in different stimulus dimensions is always incongruent, thus any dependencies between tasks are always detrimental to processing efficacy. As discussed in Section III B, the justification for this was that, assuming independent sampling of values along different stimulus dimensions, incongruent stimuli are substantially more likely and, therefore, frequent than congruent stimuli; and that the adverse effects of interference from incongruence are usually substantially more costly than the benefits of facilitation from congruence (i.e., when the sources of information favor the same set of responses). However, there are certainly some circumstances that do not align with either or both of these assumptions. For example, some previous work has focused on circumstances in which the sources of information for two tasks are congruent [21, 83, 108], and formal modeling of such situations, both mathematical [54, 114], and neural network [27], have shown how this can lead to facilitation, sometimes referred to as “super capacity”). It seems to expect that an elaboration of the framework presented here, that takes into account both the relative benefits of congruence versus costs of incongruence, and the likelihood of each, should be able to explain the effects of both. However, this remains an important direction for future research.

### Task availability

A second simplifying assumption was that stimulus information is always available along all dimensions (i.e., relevant to all tasks), irrespective of the task to be performed. In one respect, this makes the control problem more challenging, as the only way to avoid interference from irrelevant information is through the allocation of control. However, in many realistic scenarios, information for irrelevant tasks may simply not be present in the environment, making the control problem easier by affording simpler solutions. At the same time, in other respects, assuming the information is always available for all tasks also makes the control problem easier, as the system can assume that any tasks *can* be performed at any time, and the only determination that must be made is in what combinations and/or order. Again, however, in realistic environments, information may be available for some tasks but not others, posing the challenge of determining which tasks *can* be performed, before determining how to execute them. One approach to this is to consider the probability distribution of task “availability,” along the lines suggested in [19], where it is shown how reasonable assumptions about task availability may be useful as priors that shape expectations about the likelihood of opportunities for multitasking, and that favor reliance control over the investment in automaticity [19]. Further work along these lines, that explores the impact of task distribution on strategic decisions about how and when control is allocated is an important future direction, both for understanding how people manage this tradeoff and for the design of adaptive autonomous agents responsible for managing their profile of performance capabilities in complex and changing environments.

#### Complexity of tasks

Finally, as noted at the outset, the analyses presented here focused on simple “mapping” tasks, in which i) stimulus dimensions are clearly defined, ii) stimulus features along a specified input dimension are paired one-to-one with responses to be executed along a particular output dimension, and iii) processing is formulated in terms of trials in which stimulus information is presented and the expected response(s) must be generated. Needless to say, people (and machines) are capable of performing more complex tasks, i) involving environments in which stimuli do not consist of independent stimulus dimensions, ii) involving both more complex mappings from stimuli to responses and iii) more complex dynamics of execution. For example, in many cases, a task may involve the execution of actions in a particular order; and, similarly, achieving a goal may require the execution of a set of tasks in a particular order. This introduces complexity of timing and coordination that would need to be taken into account in the partitioning of tasks considered in Section IV A. Conversely, some behaviors may require coordination of the parallel execution of otherwise dissociable tasks (for example, singing and playing an instrument, or juggling). Given the large space of interactions this involves, addressing these factors in an analytically tractable form remains a challenge for future research on control and automaticity.

### D. Conclusion

Considerable progress has been made in building computational models of human cognitive performance in increasingly complex domains, both using traditional symbolic, and, more recently, neural network architectures. These have the virtue of being mechanistically explicit, and addressing empirical phenomena— concerning human performance and/or that of artificial agents—in a quantitatively precise manner. However, they remain largely descriptive, providing an account of how the human cognitive system functions, and how its processes may be implemented in the brain, or how to build machines that approximate these abilities, but without specifically addressing *why* the brain functions in those ways and/or why this would be the best way to build a machine. While a complete answer to these normative questions may never be achieved, recent work within the resource rational framework suggests that progress {emphcan be made in addressing such questions. Here, we have tried to contribute to this effort with respect to an understanding of the human capacity for cognitive control. Building on prior work using an information-theoretic approach to the allocation of control to single tasks, we have extended this to address how the system manages interactions among tasks that can arise in neural network architectures, balancing the advantages this can have through the exploitation of representational sharing for faster learning and better generalization (i.e., flexibility), against the costs it imposes by way of the need for serialization of processing (i.e. less efficient processing). The framework we have described not only undergirds a normative account of this tradeoff, and the association between capacity constraints and dependence on cognitive control, but also a normative approach to how this may be optimally balanced with the longer-term flexibility associated with the acquisition of automaticity, by modifying the task graph structure to accommodate more efficient, parallel processing through the development of separated task pathways. We hope that this approach will provide both deeper insights into the architecture of human cognition and its implementation in the brain, as well as a useful guide for how to develop artificial systems that more closely approximate the remarkable, and still uniquely human balance of flexibility and efficiency of processing.

## ACKNOWLEDGMENTS

We would like to offer a special expression of gratitude to Biswadip Dey and Kayhan Ö zcimder who helped lay some of the foundations for the work presented in this article, as well as Jonathan Pillow who helped us formulate the approach. In addition, we thank Zahra Aminzare, Adam Charles, Malory Marin, and Vaibhav Srivastava for helpful discussions. This work was supported in part by a Vannevar Bush Fellowship supported by the Office of Naval Research to JDC, and by a Schmidt Science Fellowship, in partnership with the Rhodes Trust, to SM.

## Appendix A Relationship of Probabilistic Outcome to Dynamics of Processing

In the main text we provide an information theoretic analysis of performance in the Stroop task based on the neural network described in [27]. That ignores the *dynamics* of processing, and assumes a probabilistic form for the *outcome* of processing (i.e., accuracy). This implements our assumption that the dynamics of decision processing evolve on a timescale that is much faster than the forms of control allocation and optimization that are the focus of this article (see Note V D). Here, we provide a more direct derivation of the probabilistic form we use for the outcome of processing, that relates it to dynamical systems models commonly used to address processing in simple decision making tasks (e.g., [37–39, 121]), and closely related neural network models (e.g., [27, 39]. We begin by assuming that each node in a task graph is comprised of a set of hidden units that exhibit their own dynamics of processing capable of selecting between different stimuli (see main text, Figure 1) and seek to collapse their dynamics, by focusing on steady state solutions. Inspired by the formalisms presented in [37] for two alternative forced choice decisions, and generalized to multi-choice decisions in [121], we can consider each set of processing hidden units in a population as making a decision at that layer of the network, and hidden node (*H*_*x*_) within that population as representing a choice alternative at that level of processing, which accumulates evidence for that choice *y*_*x*_ with rate *ω*_*x*1_. Following [37], we can estimate the likelihood of activating *R*_1_ in response to the hidden unit *H*_*x*_ that provides its input as:

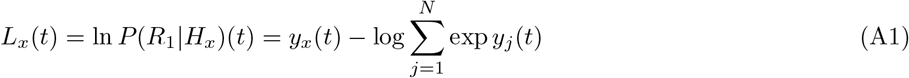

which amounts to

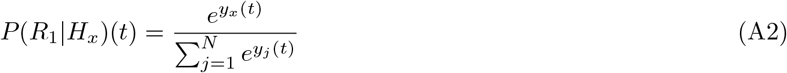

If evidence accrues with rates ***ω***, at time *t* the evidence accrued will be *t****ω***, which is a constant factor for all ***ω***, and can thus be renormalized away, implicitly defining the system’s natural temporal scale. Accordingly, the probability that the response is driven by node *x* can be expressed as :

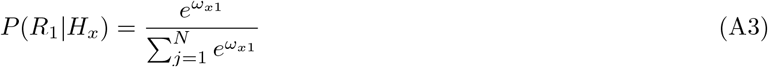

Finally, for each response node (representing the set of responses within a given response dimension), the possibility of a *no-response choice* outcome can be accommodated by adding a dummy node *H*_0_. For consistency with the previous sections, we will call the contribution to this term log 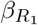, corresponding in effect to self-inhibition of units in the response node. Thus, in aggregate, we have the set of equations

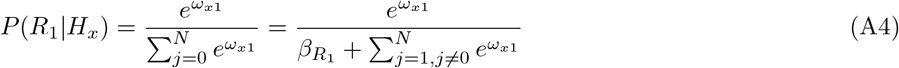

that satisfy the closure property 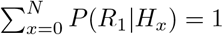. These are the equations used in the main body of the text, for describing the behavior of nodes in a task graph. In the Discussion, we consider whether and how the assumptions on which this simplification is based impact finer grained reward rate computations that take account of the dynamics of processing, and the corresponding allocation of control.

## Appendix B Full explanation of efficiencies for task architectures of Figure 5

List of extended multitasks 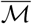 for the examples in Figure 5

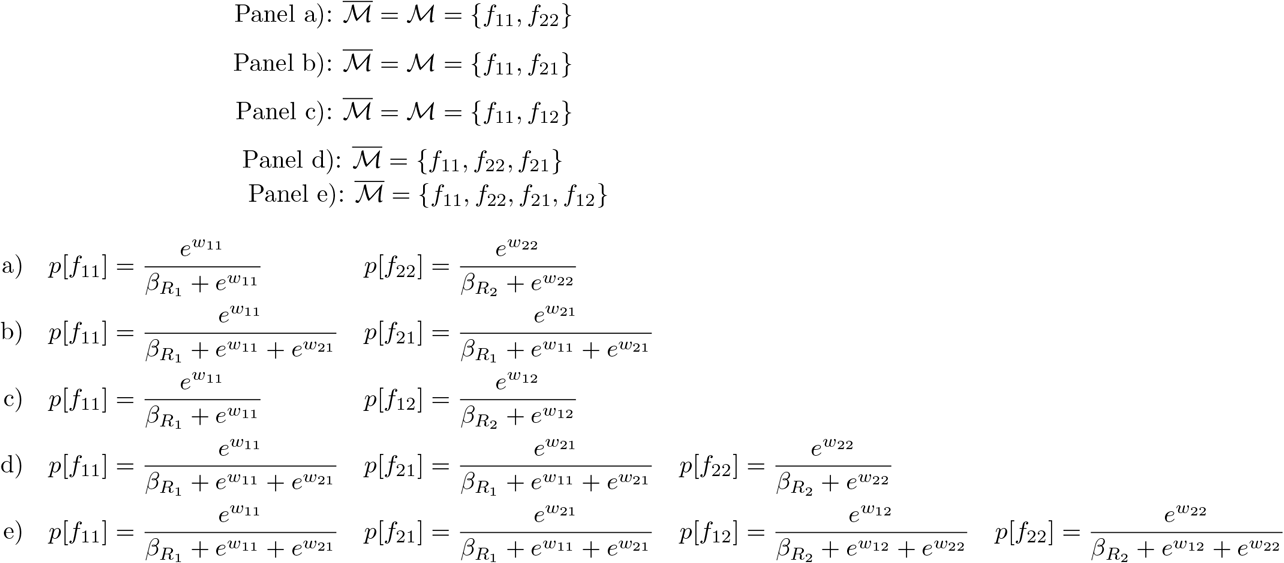

**Table I.** Edgelist for the Stroop task network in Figure 4. Here *ω*_*w*_ = 1.5, *ω*_*m*_ = 2, *ω*_*s*_ = 3, and *κ* = 1.2. For each different realization, we add .5 ∗ *ϵ* with *ϵ* a uniformly sampled noise in (0, 1). The *β* values were all set to 3 in this case.

| edge | weight w/o noise |
| --- | --- |
| 0 ($S_1^1, H_1^1$) | $\omega_m$ |
| 1 ($S_1^1, H_1^2$) | $-\omega_w$ |
| 2 ($S_1^2, H_1^2$) | $\omega_m$ |
| 3 ($S_1^2, H_1^1$) | $-\omega_w$ |
| 4 ($S_2^1, H_2^1$) | $\kappa * \omega_s$ |
| 5 ($S_2^1, H_2^2$) | $-\kappa * \omega_w$ |
| 6 ($S_2^2, H_2^2$) | $\kappa * \omega_s$ |
| 7 ($S_2^2, H_2^1$) | $-\kappa * \omega_w$ |
| 8 ($H_1^1, R^1$) | $\omega_m$ |
| 9 ($H_1^1, R^2$) | $-\omega_w$ |
| 10 ($H_1^2, R^2$) | $\omega_m$ |
| 11 ($H_1^2, R^1$) | $-\omega_w$ |
| 12 ($H_2^1, R^1$) | $\kappa_s^\omega$ |
| 13 ($H_2^1, R^2$) | $-\kappa * \omega_m$ |
| 14 ($H_2^2, R^2$) | $\kappa_s^\omega$ |
| 15 ($H_2^2, R^1$) | $-\kappa * \omega_m$ |

**Table II.** Edge weights for the graph in Figure 14. Here *ω*_*w*_ = 1.5, *ω*_*s*_ = 6, and *κ* = 1.8. For each different realization, we add .5 ∗ *ϵ* with *ϵ* a uniformly sampled noise in (0, 1).

| edge | weight w/o noise |
| --- | --- |
| 0 (S1, H1) | $\omega_s$ |
| 1 (H1, R1) | $\omega_s$ |
| 2 (S2, H2) | $\kappa * \omega_s$ |
| 3 (S2, H3) | $\omega_w$ |
| 4 (H2, R1) | $\omega_s$ |
| 5 (H2, R2) | $\kappa * \omega_s$ |
| 6 (H3, R2) | $\omega_w$ |

## Appendix C: Tables

